# Endo-Lysosome-Targeted Nanoparticle Delivery of Antiviral Therapy for Coronavirus Infections

**DOI:** 10.1101/2023.05.08.539898

**Authors:** Anton Petcherski, Brett M Tingley, Andrew Martin, Sarah Adams, Alexandra J Brownstein, Ross A Steinberg, Byourak Shabane, Jennifer Ngo, Corey Osto, Gustavo Garcia, Michaela Veliova, Vaithilingaraja Arumugaswami, Aaron H Colby, Orian S Shirihai, Mark W Grinstaff

**Affiliations:** Department of Medicine, Division of Endocrinology, David Geffen School of Medicine, University of California Los Angeles, Los Angeles, CA, USA; Department of Biomedical Engineering, Boston University, Boston, MA, USA; Molecular Cellular Integrative Physiology, University of California Los Angeles, Los Angeles, CA, USA; Department of Molecular and Medical Pharmacology, David Geffen School of Medicine, University of California Los Angeles, Los Angeles, CA, USA; Department of Chemistry, Boston University, Boston, MA, USA

**Author notes:** co-first (Anton Petcherski, Brett M Tingley).

**Keywords:** COVID-19, SARS-CoV-2, mefloquine, nanoparticles, lysosomes, endocytosis

## Abstract

SARS-CoV-2 can infect cells through endocytic uptake, a process which is targeted by inhibition of lysosomal proteases. However, clinically this approach to treat viral infections has afforded mixed results, with some studies detailing an oral regimen of hydroxychloroquine accompanied by significant off-target toxicities. We rationalized that an organelle-targeted approach will avoid toxicity while increasing the concentration of the drug at the target. Here we describe a lysosome-targeted, mefloquine-loaded poly(glycerol monostearate-co-ε-caprolactone) nanoparticle (MFQ-NP) for pulmonary delivery via inhalation. Mefloquine is a more effective inhibitor of viral endocytosis than hydroxychloroquine in cellular models of COVID-19. MFQ-NPs are less toxic than molecular mefloquine, 100-150 nm in diameter, and possess a negative surface charge which facilitates uptake via endocytosis allowing inhibition of lysosomal proteases. MFQ-NPs inhibit coronavirus infection in mouse MHV-A59 and human OC43 coronavirus model systems and inhibit SARS-CoV-2-WA1 and its Omicron variant in a human lung epithelium model. This study demonstrates that organelle-targeted delivery is an effective means to inhibit viral infection.

## INTRODUCTION

To date, severe acute respiratory syndrome coronavirus 2 (SARS-CoV-2) has infected over 763 million individuals resulting in over 6.9 million deaths globally, causing significant harm to public health and resulting in a sizable humanitarian and socioeconomic burden (Nicola *et al*, 2020; WHO, 2023). To combat this crisis, Pfizer and Moderna, among others, developed highly efficacious vaccines (BNT162b2 and mRNA-1273, respectively) at unprecedented speeds which display >94% efficacy in preventing COVID-19 illness, including severe disease (Baden *et al*, 2021; Polack *et al*, 2020). Despite rapid development and distribution of vaccines, the virus continues to gain genetic mutations in regions that have been targeted by recently developed prophylactics and treatments. Consequently, neutralization-resistant SARS-CoV-2 variants continue to spike worldwide case numbers and mortality (Dong *et al*, 2020; Garcia-Beltran *et al*, 2021).

Concurrently with vaccine development, multiple clinically approved drugs were rapidly repurposed to treat COVID-19 patients (Serafin *et al*, 2020). These include antimalarial agents such as chloroquine/hydroxychloroquine (CQ/HCQ), protease inhibitors such as lopinavir/ritonavir (LPV/RTV), and viral transcription inhibitors such as remdesivir (RDV). Despite encouraging *in vitro* reports, the clinical use of CQ/HCQ and LPV/RTV for patients with COVID-19 resulted in minimal or no clinical benefit over the standard of care (Cao *et al*, 2020; Group, 2020; Horby *et al*, 2020; Patel *et al*, 2021; WHO, 2021). Viral transcription inhibitors such as RDV and other RNA dependent RNA polymerase (RdRp) inhibitors block the viral transcription machinery from proceeding by inhibiting nucleotide incorporation into the viral mRNA (Malone *et al*, 2021; Shannon *et al*, 2022). Clinical trials evaluating RDV afforded mixed outcomes wherein RDV was superior to placebo in shortening the time to recovery in adults who were hospitalized with mild-to-severe COVID-19 in three randomized, controlled clinical trials (Beigel *et al*, 2020; Goldman *et al*, 2020; Spinner *et al*, 2020) but failed to demonstrate improved clinical outcomes as indicated by mortality rates, initiation of ventilation, and total duration of hospital stay in both the WHO Solidarity trial and DisCoVeRy trial (Ader *et al*, 2022; WHO, 2021). Emerging evidence suggests that RDV improves clinical outcomes, but only if administered within an early time window after infection (Gottlieb *et al*, 2022; Heil & Kottilil, 2022).

In addition to the early use of repurposed drugs, other small molecule viral inhibitors were developed such as nirmatrelvir (Pfizer, Paxlovid) and Molnupiravir (Merck), which received Food and Drug Administration (FDA) approval a year after RDV. Nirmatrelvir is an orally bioavailable viral 3CL^pro^ inhibitor used in combination with ritonavir to slow the metabolism of the drug (Owen *et al*, 2021). In a randomized, controlled trial for unvaccinated, non-hospitalized adults at high risk for progression to severe COVID-19, Paxlovid decreased the risk of progression to severe symptoms by 89% over placebo as determined by hospitalization rate and mortality (Hammond *et al*, 2022). More recently, Pfizer terminated the EPIC-SR trial (NCT05011513) as Paxlovid displayed no clinical benefit over placebo with regard to COVID-19 symptom relief in non-hospitalized symptomatic adults who are at low risk of progressing to severe illness. Additionally, further *in vitro* testing in both a live SARS-CoV-2 and vesicular stomatitis virus (VSV)-based pseudo-virus model suggest that selective pressure may lead to 3CL^pro^ mutations conferring nirmatrelvir resistance to new viral mutants (Heilmann *et al*, 2022; Jochmans *et al*, 2022). Molnupiravir is a prodrug nucleoside analog which causes the accumulation of significant point mutations in replicated viral transcripts (Kabinger *et al*, 2021). In a randomized, controlled trial for non-hospitalized, unvaccinated adults with mild-to-moderate COVID-19 symptoms, early treatment (within 5 days of symptom onset) with Molnupiravir resulted in a significant reduction in risk of hospitalization as well as mortality (Jayk Bernal *et al*, 2022). However, there is a growing concern that Molnupiravir, especially when administered at sub therapeutic doses, may result in the creation of more virulent SARS-CoV-2 mutants (Agostini *et al*, 2019). The discrepancy between experimental results and clinical outcomes for COVID-19 candidates and the potential for neutralization-resistant mutants, or creation of more virulent strains from emerging COVID-19 therapies, necessitates the development of new strategies, therapeutics, and prophylactics against COVID-19, as well as delivery systems to target the virus or host cell while minimizing off-target toxicity.

There are several small molecule drugs which were also repurposed for the treatment of coronaviruses and indicated efficacy in preclinical models, however they have not gained as much traction as CQ/HCQ and lopinavir. Nitazoxanide is a member of the thiazolide drug class, which are broad-spectrum anti-infection drugs, and shows early promise in both *in vitro* and small-scale *in vivo* trials (Blum *et al*, 2021; Mahmoud *et al*, 2020; Wang *et al*, 2020). The antiviral mechanism of action is not fully characterized, although reports suggest that nitazoxanide may target multiple stages of the SARS-CoV-2 life cycle, including endocytosis and membrane fusion, viral genome synthesis and viral protein processing, and the late stage inflammatory response (Lokhande & Devarajan, 2021). Sulfadoxine belongs to a class of drugs known to interrupt the synthesis of folic acid. Several *in vitro* reports demonstrate anti-SARS-CoV-2 activity with sulfadoxine, although it is unclear if folic acid synthesis plays a role in viral replication or if sulfadoxine has a novel unknown inhibitory action (Arshad *et al*, 2020; Touret *et al*, 2020).

Mefloquine (MFQ), a 4-quinolinemethanol similar in structure to CQ/HCQ, shows improved activity against SARS-CoV-2 over CQ/HCQ (Sacramento *et al*, 2022; Shionoya *et al*, 2021). CQ/HCQ blocks SARS-CoV-2 entry only in cells lacking transmembrane serine protease 2 (TMPRSS2), whereas expression of TMPRSS2 significantly reduces the antiviral activity of CQ/HCQ (Hoffmann *et al*, 2020; Lee *et al*, 2021). Existing literature suggests that mefloquine inhibits viral entry after viral attachment to the target cell (Sacramento *et al*., 2022; Shionoya *et al*., 2021). The exact mechanism is unknown, however mefloquine may be (1) inhibiting viral membrane fusion with the cell membrane or endolysosomal membrane, (2) inhibiting proteases responsible for processing SARS-CoV-2 S protein and exposing the fusion peptide, (3) modulating expression levels of angiotensin converting enzyme 2 (ACE2), Transmembrane protease, serine 2 (TMPRSS2), and/or cathepsin L (CTSL), or (4) promoting exocytosis of SARS-CoV-2 particles after uptake. Unlike CQ/HCQ, MFQ reduces viral load in clinically relevant cell lines, including Calu-3 and Vero E6/TMPRSS2 cells, which express both ACE2 and TMPRSS2 (Sacramento *et al*., 2022; Shionoya *et al*., 2021). Thus, MFQ is more broadly active against coronaviruses as compared to CQ/HCQ, and this dependence on the lack of TMPRSS2 may explain the discrepancy between *in vitro* and *in vivo* results reported using CQ/HCQ for the treatment of SARS-CoV-2.

Pharmacokinetic (PK) modeling and analysis further suggests that CQ/HCQ does not achieve therapeutic anti-SARS-CoV-2 concentrations *in vivo* when delivered orally (Liu *et al*, 2020; McLachlan *et al*, 1993; Touret *et al*., 2020). Moreover, multiple clinical trials assessing CQ/HCQ for the treatment of hospitalized patients with COVID-19 displayed an increased risk of drug-induced cardiac toxicities (Tleyjeh *et al*, 2021). Similar PK modeling of oral MFQ dosing predicts that plasma concentrations above the target EC_90_ can be achieved only with high doses over multiple days (e.g., 450 mg TID or 350 mg QID for 3 days) which likely leads to off-target effects (Karbwang & White, 1990; Sacramento *et al*., 2022; Shionoya *et al*., 2021). In fact, prophylactic use of mefloquine for malaria prevention is known to cause neurotoxicity/neurological adverse events (e.g., abnormal dreams, insomnia, anxiety, depressed mood, nausea, dizziness, and chronic central nervous system toxicity syndrome), however the mechanisms underlying its neurotoxic effects are poorly understood (Martins *et al*, 2021; McCarthy, 2015).

The non-specific delivery route for these small molecules (e.g., oral or intravenous) results in the administration of high doses with low drug accumulation in the target tissue (i.e., the lungs). As many of these small molecules are acutely cytotoxic beyond their therapeutic window, these high doses also lead to significant, off-target effects in tissues not infected with virus. To address the unmet need for a potent antiviral treatment for coronaviruses that locally targets the pulmonary system, we describe nanoparticles based on biocompatible components and loaded with chloroquine, mefloquine, sulfadoxine, or nitazoxanide. The particles are composed of poly(glycerol monostearate-co-ε-caprolactone) (PGC-C18), and, of these four payloads, mefloquine-loaded nanoparticles exhibit the strongest inhibitory effect on coronavirus infection. Herein, we report on negatively charged PGC-C18 nanoparticles (NPs) of 100-150 nm in diameter which physically entrap MFQ. MFQ-loaded NPs (MFQ-NPs) are rapidly taken up by cells, localize to endo-lysosomal compartments, and decrease protease activity. MFQ-NPs inhibit coronavirus infection in mouse MHV-A59 and human coronavirus OC43 model systems as well as inhibit SARS-CoV-2-WT and Omicron variant infection in a human lung epithelium cell line model.

## RESULTS

### PGC-C18 nanoparticle formation and characterization studies

First, we developed a method to prepare nanoparticles (NPs) with a spherical morphology of ∼100 nm. Polymeric NPs on this scale are amenable to delivery via inhalation and uptake by endocytosis (Geiser & Kreyling, 2010; Löndahl *et al*, 2014; Thorley *et al*, 2014). We selected poly(glycerol monostearate-co-ε-caprolactone), PGC-C18 (Fig. 1A), as it is comprised of biocompatible degradation products of glycerol, CO_2_, stearate, and 6-hydroxyhexanoic acid, and we have a large-scale GMP-compatible synthetic method for producing it, which would be useful in speeding clinical translation (Kaplan *et al*, 2016). To synthesize the polymer, we copolymerized ε-caprolactone and 5-benzyloxy-1,3-dioxan-2-one monomers via ring opening polymerization catalyzed by tin(II) 2-ethylhexanoate (Sn(Oct)_2_). We subsequently removed the benzyl-protecting groups of poly(5-benzyloxy-1,3-dioxan-2-one-co-ε-caprolactone) (PGC-Bn) via palladium-catalyzed hydrogenolysis and conjugated stearic acid to the newly exposed hydroxyl groups via a dicyclohexylcarbodiimide (DCC) coupling. Post-coupling, we confirmed the polymer structure via ^1^H NMR (Supp. Fig. 1A, B), and gel permeation chromatography (GPC) analysis reveals a molecular weight (M_n_) of 78,300 g/mol with narrow dispersity (Đ=1.67) (Supp. Fig. 1C).

**Figure 1.**
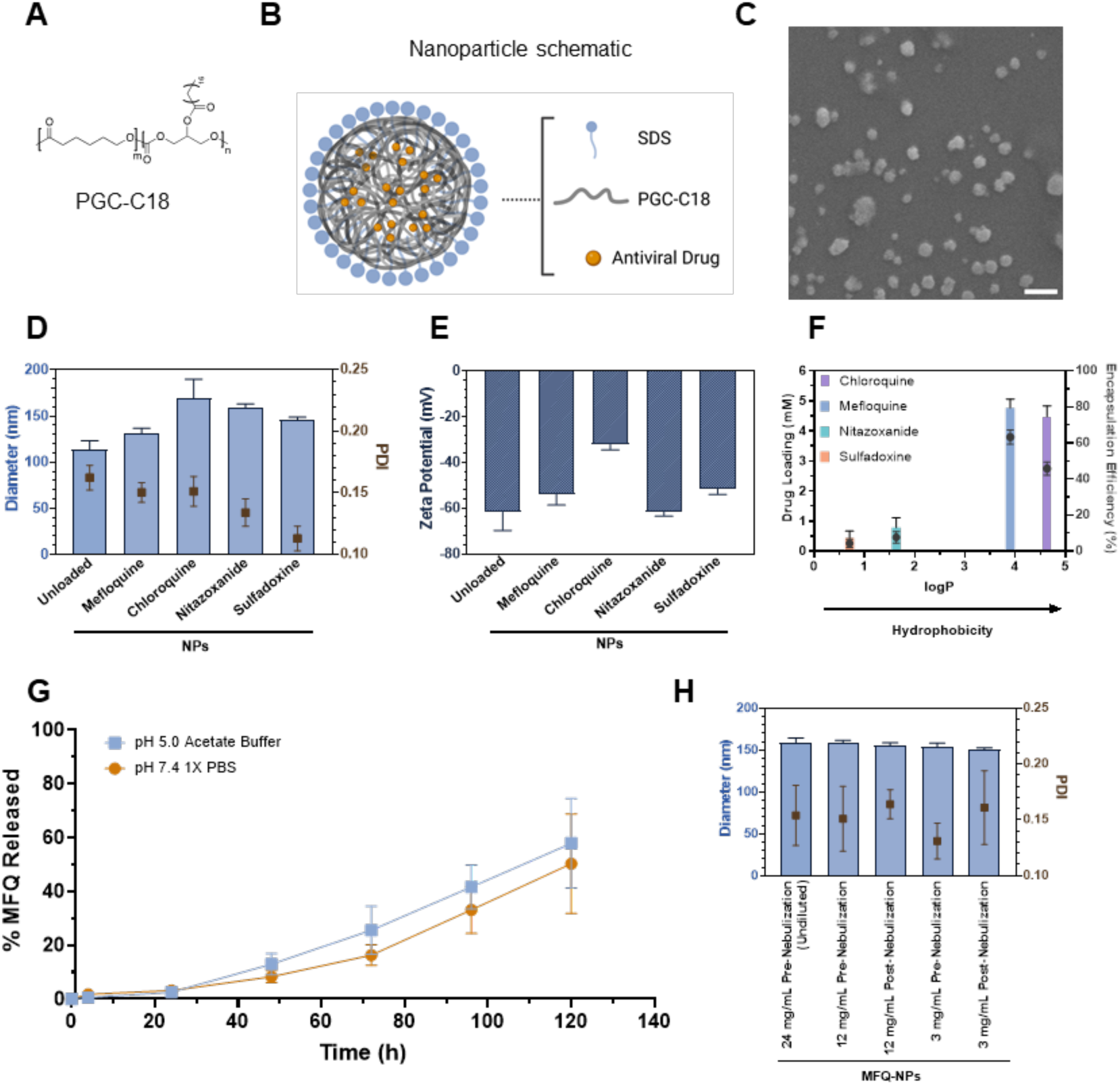
Formulation of novel PGC-C18 nanoparticles. (A) Chemical structure of poly(1,3-glycerol monostearate-co-ε-caprolactone) (PGC-C18). (B) Schematic structure of PGC nanoparticle containing sodium dodecyl sulfate surfactant and antiviral drug payload. (C) Electron micrographs of unloaded PGC-NPs demonstrate sizes around 100 nm and round morphology. Scale Bar=200 nm. (D) NP size and polydispersity measurements using dynamic light scattering (DLS) confirm nanoparticle sizes around 100-150 nm for unloaded and small molecule loaded NPs. (E) Charge measurement of NPs using DLS. (F) Encapsulation efficiency of drug compounds in NPs as measured by high performance liquid chromatography (HPLC). Drug loading concentrations are graphed as colored bars and measured on the left y-axis. Compound encapsulation efficiencies are graphed as dots (●) and measured on the right y-axis. (G) Mefloquine release from mefloquine loaded NPs (MFQ-NPs) over 5 days in pH 7.4 and pH 5.0 release buffer. Release is plotted as a % of drug released relative to initial drug loading at day 0. (H) Size and polydispersity measurements by DLS confirm nanoparticle stability after nebulization with Aerogen® Solo. All experiments in A-G represent N=3-5 independent NP batches. All data are displayed as means ± SD.

We prepared drug-loaded NPs via the solvent evaporation method (Ekladious *et al*, 2017) using sodium dodecyl sulfate as the surfactant to stabilize the formation of spherical nanoparticles containing a core of PGC-C18 polymer encapsulating a hydrophobic drug payload (Fig. 1B). SEM analysis of NP structure and size reveals spherical NPs of ∼100 nm diameter (Fig. 1C). Quantitative analysis using dynamic light scattering (DLS) confirms the size range of 100-150 nm with a good uniformity reflected in a polydispersity index of < 0.17 (Fig. 1D). Additionally, NP surface charge, as analyzed by DLS, is highly negative with a zeta-potential of ≤ -30 mV in unloaded and drug-loaded NPs (Fig. 1E), which imparts NP stability (Ekladious *et al*., 2017). As PGC-C18 is hydrophobic, it favors the encapsulation of hydrophobic compounds with chloroquine and mefloquine (logP values of 4.63 and 3.9, respectively) encapsulating more effectively than the less hydrophobic sulfadoxine and nitazoxanide (logP of 0.7 and 1.63, respectively) (Fig. 1F). Of the four drugs, mefloquine is the most effectively encapsulated compound with an encapsulation efficiency of ∼63%. In contrast, sulfadoxine and nitazoxanide lack sufficient encapsulation and exhibit negligible anti-viral activity *in vitro* (Supp. Fig. 2A, B). Therefore, we excluded both compounds from further studies. Likewise, chloroquine displays lower encapsulation efficiency and reduced *in vitro* activity against SARS-CoV-2 in cell models expressing TMPRSS2 compared to MFQ and, thus, we excluded it from further studies as well (Hoffmann *et al*., 2020; Sacramento *et al*., 2022). MFQ-loaded PGC-NPs (MFQ-NPs) exhibit controlled release over the span of 5 days in release buffer (i.e., pH 7.4 1X PBS or pH 5.0 Acetate buffer with 1 v/v% Tween 20) with 10-15% of loaded drug released after 48 h and 50-60% release at day 5 (Fig. 1G). MFQ-NPs display a consistent spherical morphology and narrow size distribution (Supp. Fig. 2C, D). Remarkably, MFQ-NPs retain size and dispersity after nebulization, further suggesting that this formulation is suitable for direct drug delivery into the lung (Fig. 1H).

### Nanoparticles exhibit minimal *in vitro* cytotoxicity

We evaluated NP cytotoxicity *in vitro* in HFL-1, Vero E6, and Calu-3 cell lines over 24 h via a tetrazolium-based MTS assay (Fig. 2A-C). Vero E6 African Green Monkey kidney epithelial cells and Calu-3 human lung adenocarcinoma cells are widely used models of SARS-CoV-2 infection (Hoffmann *et al*., 2020; Sacramento *et al*., 2022), whereas HFL-1 cells are human embryonic lung fibroblasts. Unloaded (empty) NPs are relatively non-cytotoxic until dosed at high concentrations (> 1 mg/mL). The IC_50_ values for MFQ-NPs are 42, 54, and 135 μg/mL (corresponding to approximately 7.3, 9.4, and 23.5 µM of loaded MFQ) in HFL1, Calu-3, and Vero E6 cells, respectively. For reference, the IC_50_ values for MFQ in DMSO are 11.3, 12.3, and 16.6 μM for HFL1, Calu-3, and Vero E6 cells, respectively. The vehicle itself (i.e., DMSO) is not cytotoxic at equivalent concentrations without MFQ (Supp. Fig. 2E). Notably, MFQ-NP treatments over 72 h show reduced cytoxicity compared to the 24 h timepoints in Vero E6 and Calu-3 cells when assessed with the CellTiter Blue assay (Fig. 2D-F). For the 72 h timepoints, MFQ-NP IC_50_ values are 206, and 269 μg/mL (corresponding to approximately 35.8, and 46.8 µM of loaded MFQ) for Calu-3, and Vero E6 cells, respectively. For MFQ in DMSO, the IC_50_ values are 18.8 and 19.6 μM for Calu-3 and Vero E6 cells respectively, using this assay. These data suggest that MFQ-NPs mitigate MFQ cytotoxicity via slowing the release of MFQ into the cytosol compared to the “bolus” kinetics of free drug dosing.

**Figure 2.**
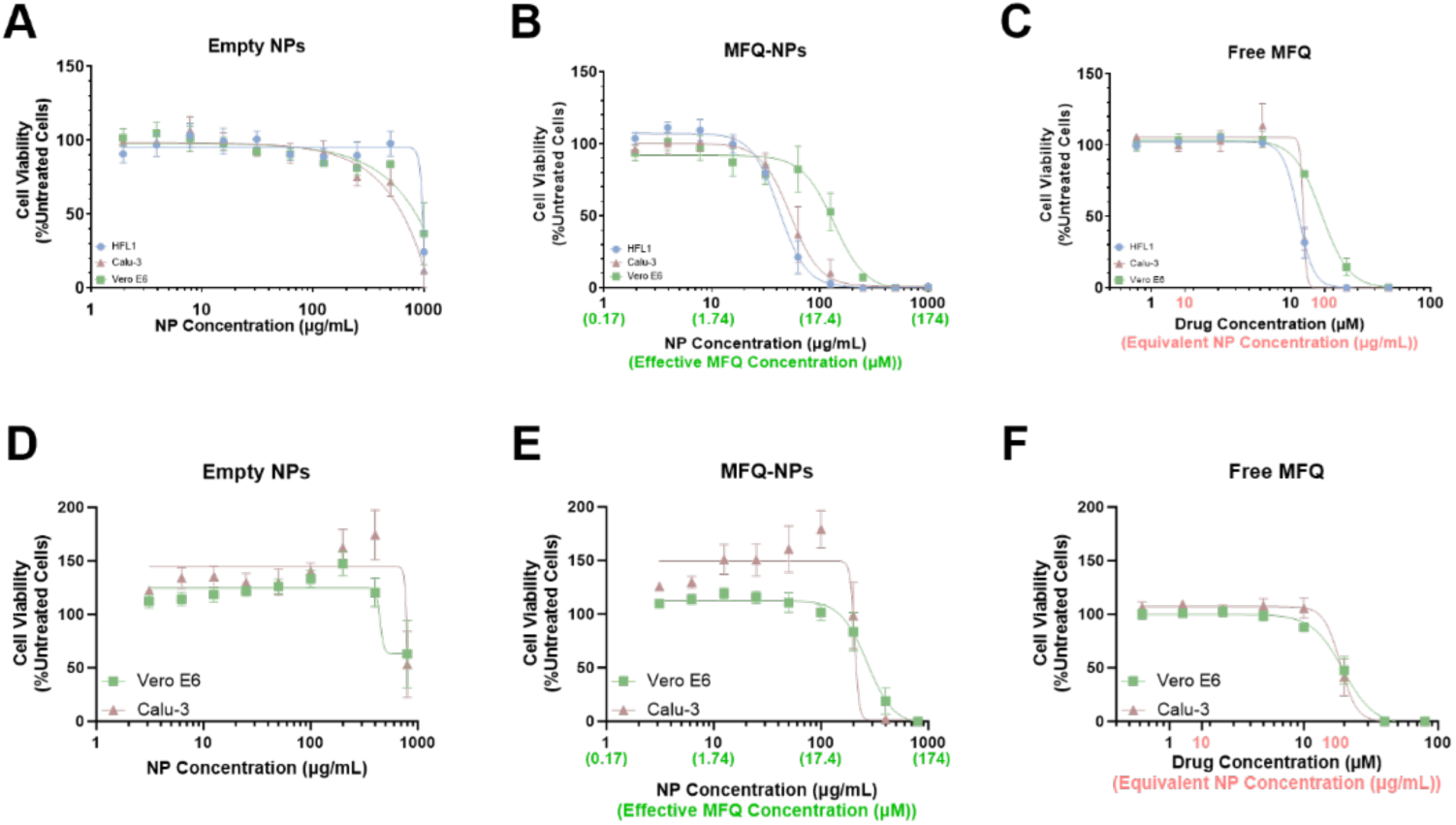
Cytotoxicity of PGC-C18 nanoparticles. Cell viability measurements by MTS assay in HFL1, Calu-3, or Vero E6 cells treated with (A) unloaded PGC-NPs, (B) Mefloquine loaded NPs (MFQ-NPs), (C) free molecular MFQ for 24 h or by CellTiter Blue assay in Calu-3 or Vero E6 cells treated with (D) unloaded PGC-NPs, (E) MFQ-NPs, (F) free MFQ for 72 h. All data are displayed as means ± SD. Each independent experiment represents the mean of n=3 technical replicates. All experiments represent N=3-6 biological replicates.

### PGC-NPs target the lysosome

To observe NP uptake, we formulated an NP containing covalently linked rhodamine B fluorophore (Rho-NPs) for analysis by flow cytometry and fluorescence microscopy. Flow cytometry reveals a rapid increase in rhodamine fluorescence after as little as 5 min of Rho-NP incubation with Calu-3, Vero E6 and HFL1 cells that steadily increases up to 24 h of Rho-NP incubation (Fig. 3A-B, Supp. Fig. 3A-E).

**Figure 3.**
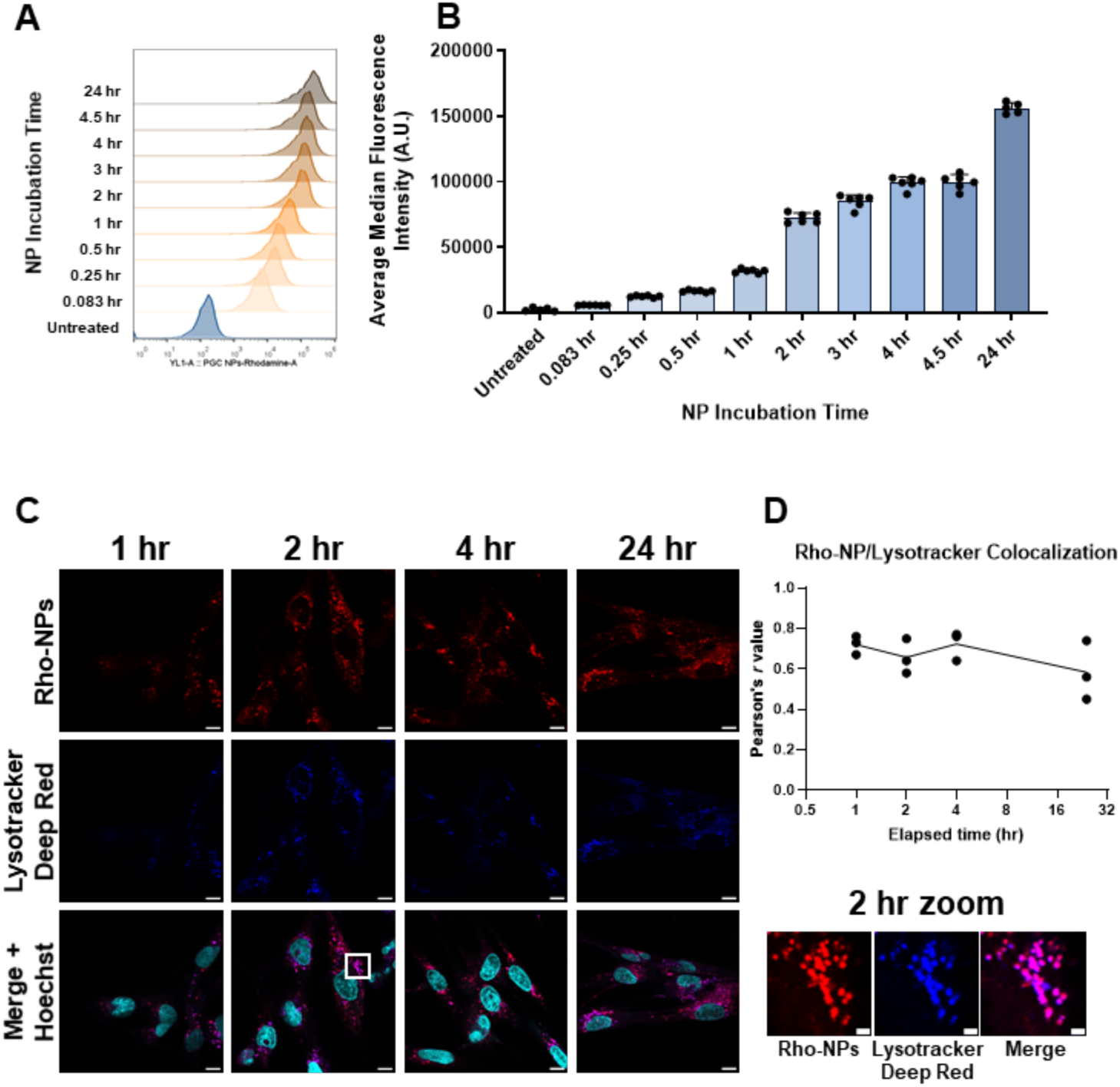
Nanoparticle uptake and localization. Rho-NP uptake measured by flow cytometry in Calu-3 cells and displayed as (A) representative intensity histogram or (B) median intensity bar graph. (C) Confocal microscopy imaging of Rho-NPs and lysosomes labelled with Lysotracker Deep Red in HFL1 cells after 1 h, 2 h, 4 h, and 24 h of NP incubation demonstrates very high levels of colocalization (quantified as Pearson’s *r* coefficient in (D)). Scale bars=10 µm; zoom-in box scale bars=2 µm. All experiments represent N=3 biological replicates which represent the mean of at least n=3 technical replicates. Intensity histogram displays a single representative cell population per timepoint. Intensity bar graph is displayed as medians ±SEM. The imaging experiment represents ∼50 cells per timepoint. Pearson’s coefficient was calculated from 3 individual images and is displayed as means ± SEM.

As flow cytometry by itself is insufficient to demonstrate Rho-NP internalization rather than just surface adsorption, we performed confocal fluorescence microscopy to confirm localization of Rho-NPs in the endo-lysosomal system after cellular uptake (Fig. 3C, D). Indeed, after 1 h of Rho-NP incubation, the majority of Rho-NP fluorescence signal colocalizes with the lysosomal live-cell dye Lysotracker Deep Red (Pearson’s coefficient, *r* = 0.74). Rho-NP colocalization with Lysotracker is consistent over a period of 24 h (Fig 3C, D).

### MFQ-PGC-NPs inhibit lysosomal activity

Using Vero E6 cells, we evaluated changes in lysosomal pH with the ratiometric probe Lysosensor-Dextran Yellow/Blue. Free MFQ and MFQ-NPs do not increase lysosomal pH but, rather, induce further acidification (Fig. 4A, B). Lysosomal pH of cells treated for 24 h with free MFQ or MFQ-NPs decreases from pH 5.1 to 4.4 (p<0.05). In contrast, treatment for 2 h with 200 nM bafilomycin A1 increases lysosomal pH to 5.7 (p<0.01). Unloaded NPs exert no effect on lysosomal pH. Although MFQ-NPs increase lysosomal acidity, this effect does not correlate with an increase in lysosomal function as free mefloquine and MFQ-NPs promote lysosomal accumulation in Calu-3 and Vero E6 cells (Fig. 4C, D, Supp. Fig. 3F, G). Notably, 24 h treatments with free MFQ and MFQ-NPs afford a 2.5-3-fold increase in the amount of lysosomal staining area in Calu-3 cells (p<0.001) and a 6.2-6.9-fold increase in Vero E6 cells (p<0.0001). Bafilomycin A1 treatment strongly inhibits Lysotracker staining presumably by dissipating lysosomal pH. Free MFQ and MFQ-NP treatments, at concentrations above 15 µM, reduce lysosomal protease activity to the same degree as treatment with bafilomycin A1 or the lysosomal protease inhibitors pepstatin A and E64d by 57-75% (p<0.01) (Fig. 4E). Interestingly, MFQ concentrations below 10 µM increase lysosomal protease activity by up to 47% (p<0.01), potentially due to the activation of compensatory lysosomal acidification. Unloaded NPs exhibit no effect on lysosomal protease activity.

**Figure 4.**
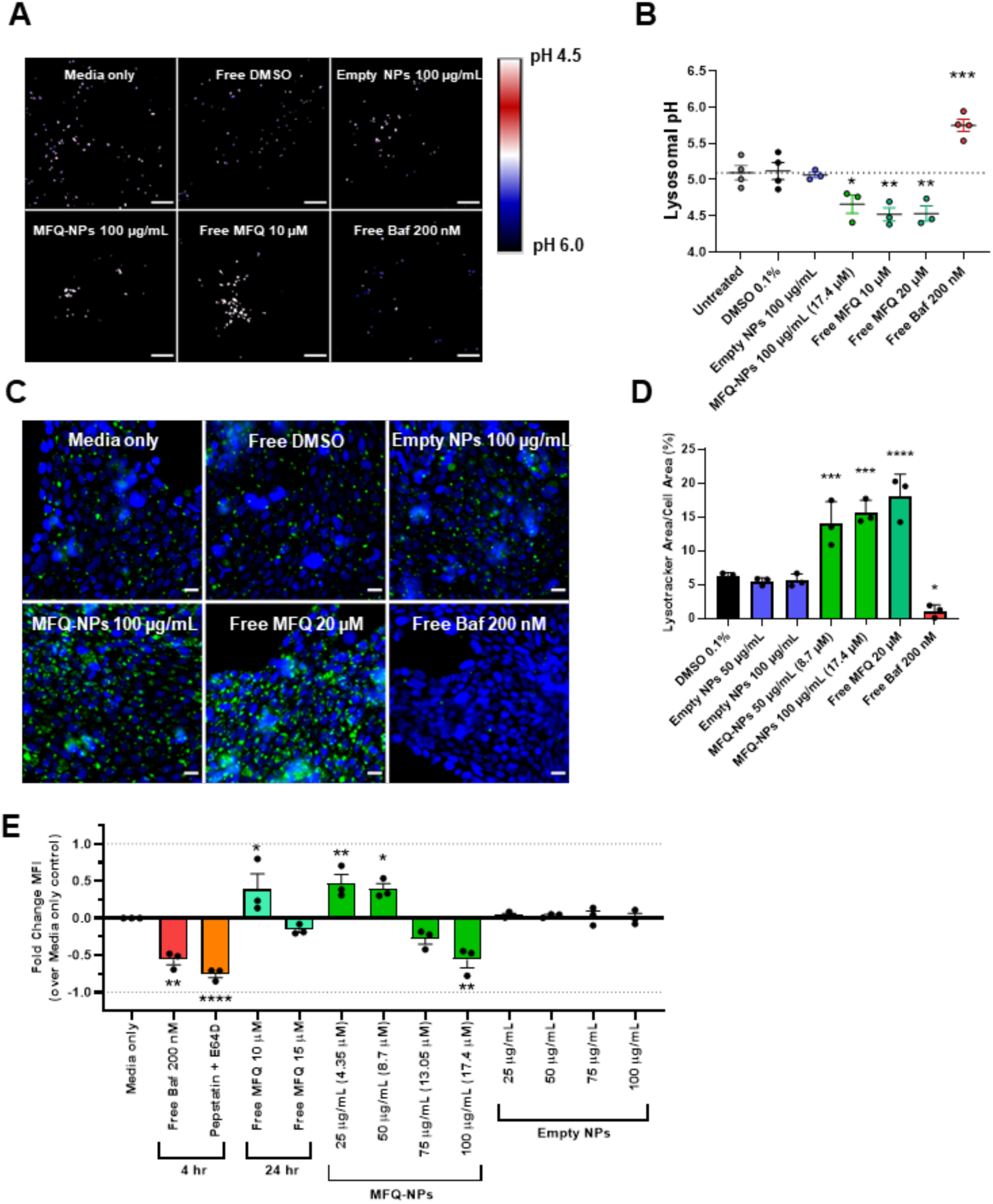
Nanoparticle effects on lysosomal pH and protease activity. (A) Representative confocal images of lysosomal pH measurements using Lysosensor-Dextran Yellow/Blue in NP (±MFQ) 100 µg/mL, free MFQ 10 µM, bafilomycin 200 nM, or control treated Vero E6 cells. Scale bars=10 μm. (B) Quantification of lysosomal pH. (C) Representative images of Calu-3 cells treated with NPs (±MFQ) 100 µg/mL, free MFQ 20 µM, bafilomycin 200 nM, or control and stained with Hoechst 33342 (blue) and Lysotracker Green (green) to probe for lysosome accumulation. Scale bars=20 μm. (D) Quantification of lysosomal accumulation. (E) Quantification of lysosomal protease activity by DQ-Red BSA assay in Vero E6 cells treated with NPs (±MFQ), free MFQ, bafilomycin A1, pepstatin A + E64d, or controls at the indicated concentrations. Statistical significance was determined by one-way ANOVA: *p<0.05, **p<0.01, ***p<0.001, ****p<0.0001 against untreated controls. Each independent Lysosensor imaging experiment represents the mean of n=10-20 individual images. All other biological replicates represent the mean of n=3 technical replicates. All experiments represent N=3 biological replicates. All data are displayed as means ± SEM.

### MFQ-PGC-NPs inhibit murine coronavirus MHV-A59 infection

Mouse hepatitis virus A59 (MHV-A59) is a beta coronavirus that infects mice displaying neuro-, hepato-, and pneumotropism depending on the route of infection (Cowley & Weiss, 2010). Like SARS-CoV-2, MHV-A59 displays multi-organ involvement and leads to more severe pneumonia in aged individuals (Ryu *et al*, 2021). L929 mouse fibroblasts infected with MHV-A59-GFP produce a lytic infection with extensive syncytia formation (Supp. Fig. 4A-C). Preliminary studies using L929 cells indicated that MHV-A59-GFP infection, assessed by GFP-positivity, peaked at 20-24 h post infection, after which cell death, measured by Annexin-V-positivity, rapidly sets in (Supp. Fig. 4A, B). Since dying cells rapidly lose their GFP+ signal, we included syncytia formation as an additional measure of viral infection frequency (Supp. Fig. 4C). We adopted a preincubation protocol that allowed us to observe MFQ-NP effects on viral binding and uptake (Fig. 5A). Pre-incubation with empty PGC-NPs exerts no preventive effect on MHV-A59-GFP viral infection (Fig. 5B, C). In contrast, pre-incubation with MFQ-NPs and free MFQ reduces the amount of GFP+ cells and the incidence of syncytia formation in a dose-dependent manner (Fig. 5B-E) by 29% at 12.5 μg/mL and up to 97% at the highest concentration of 100 μg/mL (p<0.0001). At the highest tested concentrations (100 μg/mL for MFQ-NPs and 10 μM for free MFQ) a slight reduction in overall cell counts occurs compared to infected controls (∼20-27% reduction, p<0.05). As a positive control, Remdesivir completely inhibits viral replication at a concentration of 10 μM. Occasionally, individual cells treated with MFQ or MFQ-NPs still exhibit GFP-positive fluorescence; however, the infection did not spread in the culture (Fig. 5B).

**Figure 5.**
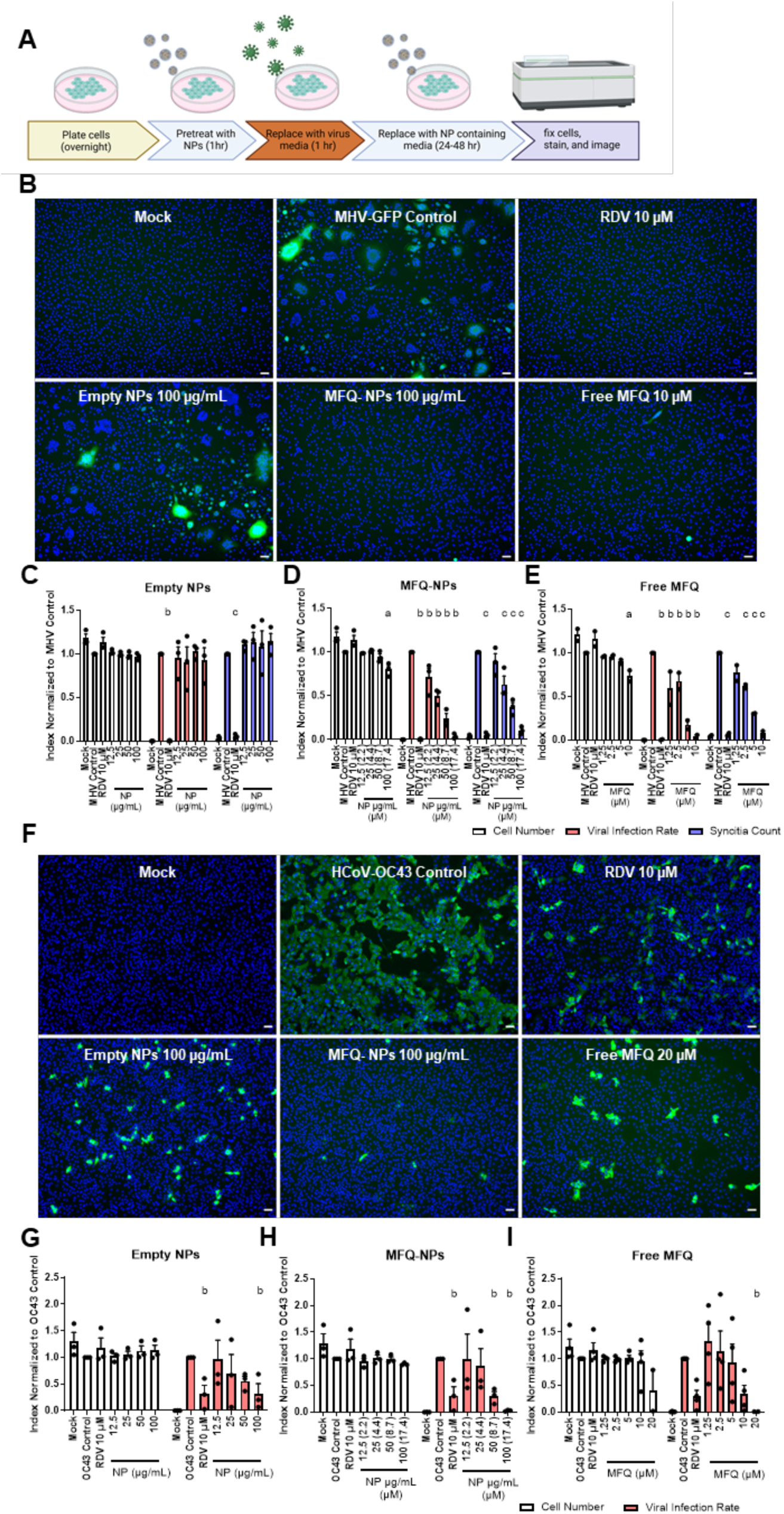
PGC-NPs loaded with MFQ are effective at inhibiting MHV and HCoV-OC43 infection. (A) Schematic describing the treatment and infection sequence of prophylactic NP treated MHV-GFP or HCoV-OC43 infected cells. (B) Representative images of L929 cells pre-treated with control, remdesivir (RDV, 10 µM), free MFQ (10 µM), NPs (±MFQ) (100 µg/mL) and infected with MHV-GFP (MOI 0.1) for 24 h. Nuclei were stained with DAPI (blue), and virus infected cells were visualized by GFP-positive signal (green) or syncytia formation. Scale bars=50 μm. Quantification of cell numbers, GFP-area per image, and syncytia in unloaded NPs (C), MFQ-NPs (D) or free MFQ (E) treated cells. Statistical significance was determined by two-way ANOVA: a (at least) p<0.05 against MHV-Control cell numbers; b (at least) p<0.05 against MHV-Control infected cell area; c p<0.05 against MHV-Control syncytia count. (F) Representative images of Vero E6 cells pre-treated with control, RDV (10 µM), free MFQ (20 µM), NPs (±MFQ) (100 µg/mL) and infected with HCoV-OC43 (MOI 1) for 48 h. Nuclei were stained with DAPI (blue), and virus infected cells were visualized by immunofluorescence staining against pan coronavirus nucleocapsid (green). Scale bars=50 μm. (D) Quantification of cell numbers and fractions of infected cells in empty NP (G), MFQ-NP (H) or free MFQ (I) treated cells. Statistical significance was determined by two-way ANOVA: a (at least) p<0.05 against OC43-Control cell numbers; b (at least) p<0.05 against OC43-Control infected cell fraction. All biological replicates represent the mean of n=3 technical replicates. All experiments represent N=3 biological replicates. All data are displayed as means ± SEM.

### MFQ-PGC-NPs inhibit human coronavirus OC43 infection

Like SARS-CoV-2, HCoV-OC43 is a beta coronavirus that causes respiratory tract infections in humans; however, HCoV-OC43 infections are generally mild cold-like symptoms. Vero E6 cells, infected with a high titer of HCoV-OC43 (MOI=1), exhibit pronounced virus positivity after 48 h (Fig. 5F, ‘Virus infected Control’). Interestingly, remdesivir treatment results in an incomplete protection against viral infection at 10 µM, reducing viral infection rates by about 69% (p<0.05). Surprisingly, empty NPs inhibit viral infection at the highest tested concentration of 100 μg/mL with inhibition similar in magnitude to remdesivir controls (68% reduction, p<0.05) (Fig. 5F, G). MFQ-NPs display a concentration dependent inhibition of viral infection that is statistically significant at 50 μg/mL (70% reduction, p<0.05) (Fig. 5F-I) and reaches almost 100% at 100 μg/mL, whereas free MFQ inhibits viral replication at 10 μM but is only significant at 20 μM, at which concentration one observes substantial cytotoxicity.

### MFQ-PGC-NPs inhibit infection with SARS-CoV-2 WT-WA1 and Omicron BA.1 variants

To assess MFQ-NP efficacy against the COVID-19 pandemic virus, SARS-CoV-2, we utilized two different cell lines. Vero E6 cells that do not express human TMPRSS2, therefore favoring SARS-CoV-2 infection by endocytosis, and Calu-3 human alveolar epithelial adenocarcinoma cells with a high expression of TMPRSS2, thus favoring SARS-CoV-2 spike cleavage and fusion at the plasma membrane. We initially adopted a similar pre-infection treatment which did not include washing-off the viral inoculate (Fig. 6A). In Vero cells, remdesivir (10 μM) effectively inhibits SARS-CoV-2 infection (98% reduction, p<000.1) (Supp. Fig. 5A, B), whereas empty PGC-NPs exhibit no effect, and MFQ-NPs reduce infection only at 100 μg/mL (79% reduction, p<0.0001). Moreover, free MFQ reduces infection only at 20 μM (93% reduction, p<0.001) without exhibiting significant toxicity (Supp. Fig. 5A-D). In Calu-3 cells, MFQ-NP treatment at 100 μg/mL significantly reduces the number of SARS-CoV-2 positive cells (91%, p<0.05) and similarly, strong inhibition was observed by free MFQ at 20 μM (92% reduction, p<0.05) (Fig. 6B-E). In addition to the ancestral SARS-CoV-2 variant, WT-WA1, we assessed MFQ-NP efficacy against Omicron BA.1. After preincubation, empty PGC-NPs exert a small inhibitory effect on Omicron infection at 25-100 μg/mL (26-43% reduction, p<0.01-0.0001) (Fig. 6F). MFQ-NPs significantly inhibit Omicron infection at 12.5 and 100 μg/mL (29% and 83% inhibition, p<0.05 and <0.0001 respectively) (Fig. 6G). Free MFQ significantly inhibits Omicron infection at 10 and 20 μM (60-97% inhibition, p<0.001-0.0001) (Fig. 6H).

**Figure 6.**
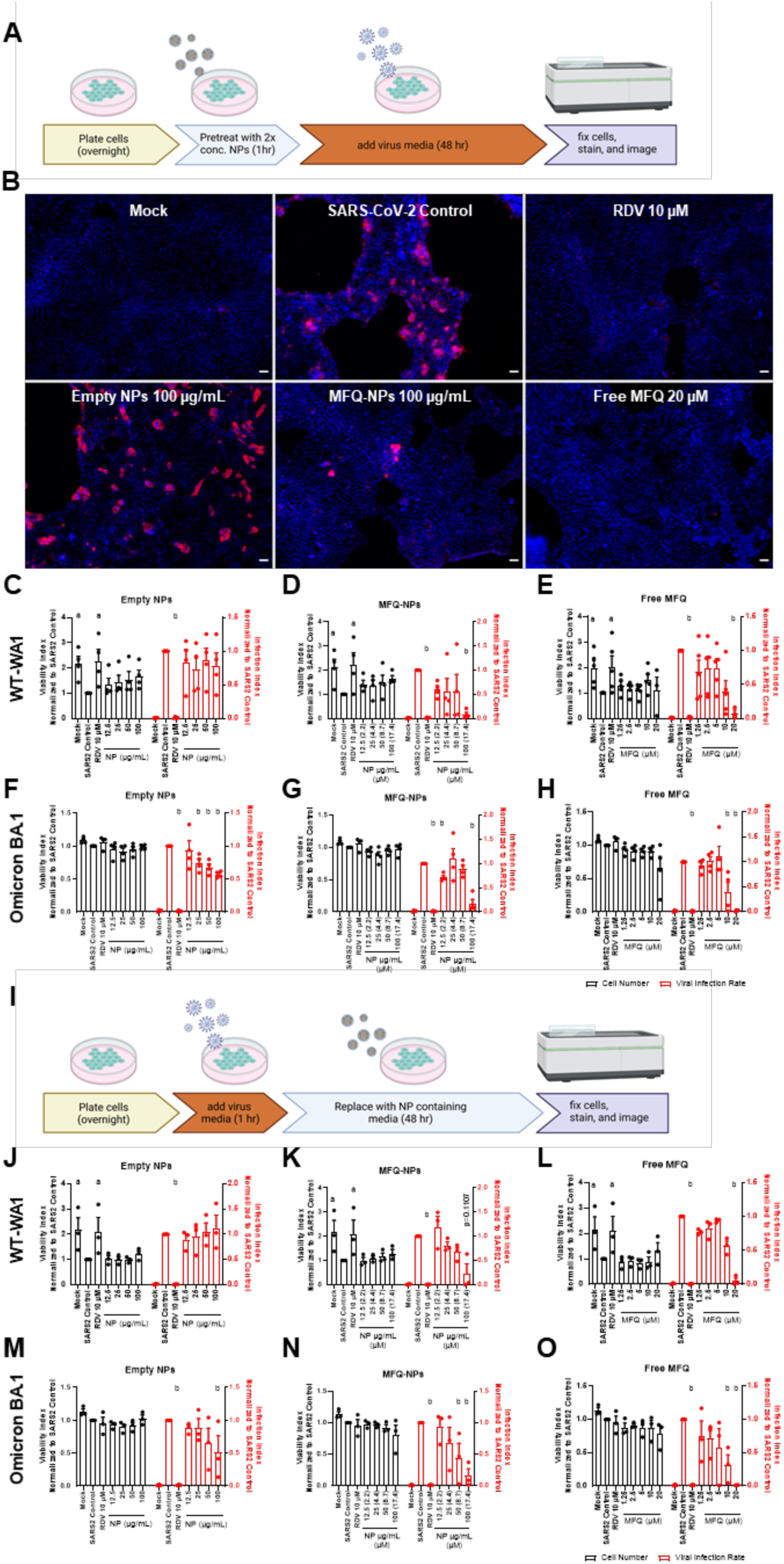
PGC-NPs loaded with MFQ are effective at inhibiting SARS-CoV-2 WT-WA1 and Omicron infection and replication. (A) Schematic describing the treatment and infection sequence of prophylactic NP treated SARS-CoV-2 infected cells. (B) Representative images of Calu-3 cells pre-treated with control, RDV (10 µM), free MFQ (20 µM), NPs (±MFQ) (100 µg/mL) and infected with SARS-CoV-2 WT-WA1 (MOI 0.2) for 48 h. Nuclei were stained with DAPI (blue) and virus infected cells visualized by immunofluorescence staining against SARS-CoV-2 N-protein (red). Scale bars=50 μm. Quantification of cell numbers and fractions of infected cells in unloaded NP (C), MFQ-NP (D) or free MFQ (E) pretreated Calu-3 cells infected with SARS-CoV-2 WT-WA1. Quantification of cell numbers and fractions of infected cells in unloaded NP (F), MFQ-NP (G) or free MFQ (H) pretreated Calu-3 cells infected with SARS-CoV-2 Omicron BA.1. Statistical significance was determined by two-way ANOVA: a (at least) p<0.05 against SARS-CoV-2 Control cell numbers; b (at least) p<0.05 against SARS-CoV-2-Control infected cell area. (I) Schematic describing the treatment and infection sequence of SARS-CoV-2 infected cells treated after infection. Quantification of cell numbers and fractions of infected cells in unloaded NP (J), MFQ-NP (K) or free MFQ (L) treated Calu-3 cells infected with SARS-CoV-2 WT-WA1. Quantification of cell numbers and fractions of infected cells in unloaded NP (M), MFQ-NP (N) or free MFQ (O) treated Calu-3 cells infected with SARS-CoV-2 Omicron BA.1. Statistical significance was determined by two-way ANOVA: a (at least) p<0.05 against SARS-CoV-2 Control cell numbers; b (at least) p<0.05 against SARS-CoV-2-Control infected cell area. All biological replicates represent the mean of n=3 technical replicates. All experiments represent N=3-4 biological replicates. All data are displayed as means ± SEM.

### MFQ-PGC-NPs inhibit post-exposure spread of SARS-CoV-2 WT-WA1 and Omicron BA.1

The experimental design used to assess MFQ-NP efficacy under prophylactic treatment conditions cannot distinguish between decreased viral binding and endocytosis, and an inhibition of viral replication. Thus, to study a potential inhibition of viral replication, we modified the protocol to include a 1 h incubation with SARS-CoV-2 inoculum before starting MFQ-NP treatments, which allows for viral attachment and uptake prior to the beginning of the treatment (Fig. 6I). Unloaded-PGC-NPs do not prevent SARS-CoV-2 WT-WA1 replication but exhibit a significant effect in Omicron infected Calu-3 cells (49% reduction, p<0.05) (Fig. 6J, M). MFQ-NPs inhibit viral replication of the ancestral WT-WA1 variant to a non-significant degree (77% reduction, p=0.11) and significantly reduce Omicron replication at 50 and 100 μg/mL (56% and 84% reduction, p<0.01 and <0.001) (Fig. 6K, N), which is similar to the result obtained with free MFQ at 10 and 20 μM (up to 98% reduction of Omicron positive cells, p<0.0001) (Fig. 6 L, O). These data suggest that MFQ-NPs inhibit viral uptake as well as replication.

### MFQ-NPs inhibit coronavirus infection in a post-attachment phase and alter expression of proteins responsible for viral uptake

We next addressed which stage in the viral life cycle free MFQ and MFQ-NPs inhibit by performing viral attachment and entry assay.s and assessing mRNA expression level of entry proteins after NP treatment.

In the case of SARS-CoV-2, the spike (S) protein binds to the ACE2 receptor on the surface of the cell. Following attachment, the viral particle is subjected to S protein cleavage by host cell proteases, either TMPRSS2 on the cell membrane or CTSL following endosomal uptake (Glowacka *et al*, 2011; Lee *et al*., 2021; Matsuyama *et al*, 2010; Zhao *et al*, 2021). S protein cleavage initiates fusion pore formation between the viral membrane and the plasma membrane or endolysosomal membrane leading to viral RNA release into the cytoplasm and viral replication. Previous work (Brufsky, 2020; Shionoya *et al*., 2021; Yao *et al*, 2022a) suggests that CQ, HCQ and MFQ may impact expression of target cell entry proteins or entry protein glycosylation. As MFQ is very similar in structure to these compounds, we sought to determine if MFQ alters expression of entry proteins responsible for SARS-CoV-2 uptake (e.g., ACE2, TMPRSS2 and CTSL) and MHV uptake (e.g., CEACAM1). To this end, Calu-3 or L929 cells were plated and allowed to adhere overnight. The following day, cell culture media was replaced with media containing free MFQ, MFQ-NPs, empty NPs, DMSO control, or untreated media for 48 h. Following treatment, RNA was isolated, and expression was measured using RT-qPCR for *hACE2*, *hTMPRSS2* and *hCTSL* in Calu-3 cells and *mCEACAM1* in L929 cells (Supp. Fig. 6A-D). We observe a dosage dependent fold-increase in expression of both *hTMPRSS2* and *hCTSL* in Calu-3 cells with free MFQ and MFQ-NPs. Conversely, the expression of *hACE2* reduces following treatment with free MFQ and MFQ-NPs. Unlike treatments containing high-dose MFQ, all other treatments (e.g., empty NPs and DMSO controls) display relatively similar expression level of entry protein transcripts compared to media only controls. In L929, the expression of *mCEACAM1* significantly decreases following high-dose MFQ treatment in free drug or NP formulations. However, empty NPs do not reduce expression of *mCEACAM1* relative to untreated control, and DMSO treated groups significantly reduced expression but not to an equal extent as the high dose (i.e., 17.4 µM) free MFQ and MFQ-NPs.

To assess whether MFQ or MFQ-NPs inhibit viral attachment, we pre-treated Vero E6 cells with varying concentrations of NPs or compounds for 1 h. Following pre-treatment, cells were exposed to SARS-CoV-2 for 1 h at 4°C to allow for viral attachment but not entry and replication (Supp. Fig. 6E). After washing away unattached virus and NPs or compounds, we extracted and quantified the viral RNA attached to the cell surface. Cell membrane associated SARS-CoV-2 RNA non-significantly reduces following treatment with free MFQ or MFQ-NPs (Supp. Fig. 6F). Surprisingly, empty NPs (p=0.007) and E64d (p=0.007) significantly inhibit viral attachment.

To evaluate whether MFQ or MFQ-NPs inhibit viral endocytosis and membrane fusion (post-attachment phase), we similarly pre-treated Vero E6 cells with varying concentrations of NPs or compounds for 1 h. Following pre-treatment, cells were exposed to SARS-CoV-2 for 1 h at 37°C to allow for viral attachment. After washing away unattached virus, we incubated the cells in growth media containing the NPs or compounds for 6 and 24 hours, respectively, before extracting and quantifying the internalized viral RNA. After only 6 h of incubation, post-attachment viral RNA significantly diminishes following treatment with free MFQ (p<0.0001), MFQ-NPs (p<0.0001) and lysosomal protease inhibitor E64d (p=0.0002) (Supp. Fig. 6G). However, empty NPs exert no effect on viral RNA levels, suggesting that MFQ is responsible for the reduction in viral RNA post-attachment. After 24 h of incubation, viral RNA significantly reduces to levels comparable to Remdesivir following treatment with free MFQ (p<0.0001), MFQ-NPs (p<0.0001) and E64d (p=0.0055). Empty NPs similarly fail to reduce viral RNA levels following a 24 h incubation (Supp. Fig 6H).

## DISCUSSION

The COVID-19 pandemic highlights the capacity of a sudden infectious disease to cause a long-lasting impact on public health. In response to the emergence of SARS-CoV-2, scientists rapidly developed prophylactic vaccines and began repurposing already availably drugs to treat hospitalized patients. In this study we describe the development of nanoparticles (NPs) using a biocompatible polymer we have developed, poly(glycerol monostearate-co-ε-caprolactone) (PGC-C18), for the pulmonary delivery of antiviral drugs, particularly for the treatment of coronavirus infection. We selected PGC-C18 due to our experience with this polymer in a different drug delivery form factor (i.e., an implantable surgical mesh), its successful completion of 10993 biocompatibility testing required by the FDA (Kaplan *et al*., 2016), and its availability via large-scale GMP production processes that would speed translation to the clinic. Furthermore, PGC-C18 exhibits superior long-term compound release properties and lacks an initial drug “burst” release associated with unmodified or short chain fatty acid modified PGC surfaces (Wolinsky et al., 2010, 2012).

Other groups have similarly leveraged polymer systems to physically encapsulate antibiotics or antimicrobials for pulmonary delivery (Al-Halifa *et al*, 2019; Coowanitwong *et al*, 2008; Ungaro *et al*, 2012). Polylactic acid (PLA) and poly(lactic-co-glycolic acid) (PLGA) are commonly employed as they are biodegradable and biocompatible polymers present in FDA and European Medicines Agency (EMA) approved products for use in humans. However, these formulations tend to form particles ≥ 200 nm with many manufacturing approaches resulting in microspheres on the µm-size scale. Multiple particle deposition studies suggest that NPs > 100 nm fail to reach deep lung tissue (i.e., the alveoli), which makes these polymer platforms a less attractive approach for prophylactics/treatments against coronaviruses, such as SARS-CoV-2, as this virus largely targets alveolar type 2 (AT2) cells, one of the major cell types that co-express ACE2 and TMPRSS2 (Geiser & Kreyling, 2010; Liu *et al*, 2021; Löndahl *et al*., 2014).

We fabricated PGC-NPs using an emulsification and solvent evaporation method which yielded NPs between 100-150 nm in diameter with low polydispersity (<0.17) and low vehicle cytotoxicity. These NPs physically encapsulate small molecule drugs, and the encapsulation efficiency increases with increasing hydrophobicity/lipophilicity (i.e., logP) of the drugs. Among the four drugs loaded - drugs chloroquine, mefloquine, sulfadoxine, and nitazoxanide - mefloquine is most effectively encapsulated compound with an encapsulation efficiency of ∼63%. MFQ-loaded NPs exhibit controlled release and retain their size and dispersity after nebulization, making them suitable for direct drug delivery into the lung. Furthermore, the NPs exert minimal *in vitro* cytotoxicity in human lung fibroblast (HFL-1), Vero E6, and Calu-3 cell lines, making them suitable for clinical translation.

Of the investigated drugs, mefloquine shows improved activity against SARS-CoV-2 over other repurposed drug candidates such as hydroxychloroquine and chloroquine yet remains under-investigated as an antiviral therapy (Sacramento *et al*., 2022; Shionoya *et al*., 2021). Jan *et al*. report that orally administered treatment of MFQ at 30 mg/kg/day for 3 days results in the absence of weight loss in SARS-CoV-2 infected hamsters (Jan *et al*, 2021). However, viral lung titers only decrease by less than one log unit and no further investigation regarding disease progression or lung histology was reported. Preliminary pharmacokinetic modeling suggests that conventional (i.e., oral) dosing of MFQ requires multiple high doses (e.g., 350 – 450 mg) daily to achieve therapeutic plasma concentration, likely impeding clinical translation of orally-dosed MFQ. Alternatively, pulmonary delivery of MFQ would enable higher concentrations in lung tissue, reduce systemic exposure and mitigate off-target toxicities (i.e., neurotoxicity), while eliminating the need for repeat daily doses. As a first step towards this goal, we designed a polymer-based nanoparticle to deliver mefloquine locally to the respiratory tract, the site of SARS-CoV-2 infection. MFQ-NPs exhibited robust and dose-dependent viral inhibition across multiple coronavirus strains including MHV, HCoV-OC43, and SARS-CoV-2 (USA-WA1/2020 and Omicron BA.1 variants). Additionally, unlike hydroxychloroquine and chloroquine, MFQ is efficacious against SARS-CoV-2 infection in cell lines expressing both ACE2 and TMPRSS2 (e.g., Calu-3) (Fig. 6 and (Sacramento *et al*., 2022; Shionoya *et al*., 2021)).

Despite mefloquine being used as an antimalarial drug similar to chloroquine, little is known about the biological response to mefloquine treatment, particularly regarding lysosomal acidification. In acute myeloid leukemia cells, mefloquine disrupts lysosomal integrity while exerting a biphasic effect on lysosomal pH (Sukhai et al., 2012; Lam Yi et al., 2019). Here, treatment with mefloquine or MFQ-NPs decreases lysosomal pH with a concomitant increase in lysotracker accumulation when compared to non-treated controls. Interestingly, further acidification as a result of mefloquine treatment corresponds to an inhibition of proteolytic degradation at high dosages. Unlike chloroquine, which is a known lysosomotropic agent that increases lysosomal pH, leading to a decrease in lysosomal proteolytic activity, mefloquine inhibits proteolytic activity while exerting the opposite effect on lysosomal pH at therapeutic dosing (Hoffmann *et al*., 2020). At sub-therapeutic dosing in our study, mefloquine treatment decreases lysosomal pH and increases proteolysis but, conversely, lysosomal accumulation increases over baseline. These effects on lysosomal pH and activity resemble a biphasic dose response (hormesis) described previously for chloroquine and its derivatives and may explain the varied outcomes in the treatment of COVID-19 patients with lysosomotropic drugs (Calabrese *et al*, 2021; Moore, 2020).

In the context of SARS-CoV-2, mefloquine is known to inhibit viral entry, specifically during post-attachment processes (Sacramento *et al*., 2022; Shionoya *et al*., 2021). Both in free drug and NP formulations mefloquine inhibits SARS-CoV-2 entry and replication to a similar degree as E64d and Remdesivir post-attachment. However, viral attachment only reduces following transient exposure to virus followed by washing. For both empty NPs and MFQ-NPs, this may be partially explained by steric hinderance of viral attachment due to nonspecific adsorption of NPs onto the cell surface. The remarkable effect of E64d inhibiting viral attachment to the cell may be explained its inhibition of secreted CTSL, which enhance coronavirus entry into cells (Zhao *et al*., 2021). To further interrogate whether mefloquine exerts any effect on the viral attachment phase, we investigated the expression level of several entry proteins responsible for SARS-CoV-2 uptake in Calu-3 cells and MHV uptake in L929 cells. In response to high dose (i.e., 17.4 µM) mefloquine in free drug or NP form, RNA transcripts for ACE2 decrease and expression increases for both TMPRSS2 and CTSL. Yao et al report similar results following treatment with hydroxychloroquine in human primary pterygium and conjunctival tissues (Yao *et al*., 2022a). There is some evidence that chloroquine and hydroxychloroquine inhibit ACE2 glycosylation (Brufsky, 2020) which may induce incorrect trafficking of ACE2 and reduce ACE2 expression. As ACE2 is an essential target cell protein for SARS-CoV-2 infection, it is plausible that prolonged (≥ 48 h) treatment with mefloquine reduces pulmonary cell ACE2 expression which leads to inhibition of viral attachment in addition to inhibition of post-attachment viral proliferation. The increase in CTSL expression may be the result of a compensatory response to the observed lysosomal accumulation and decrease in lysosomal proteolytic activity following high dose mefloquine treatments. Remarkably, treatment with hydroxychloroquine slightly increases TMPRSS2 expression levels, indicating a similar response of TMPRSS2 expression regulation to lysosomal inhibition to CTSL (Yao *et al*, 2022b). In L929 cells, murine carcinoembryonic antigen-related cell adhesion molecule (mCEACAM1) expression significantly reduces when treating with high dose free MFQ or MFQ-NPs. mCEACAM1a is an identified cognate target for MHV spike protein, thus downregulation of mCEACAM1 by mefloquine may provide a protective effect against MHV infection. Similarly, a broad reduction in cell adhesion markers including CEACAM1, 5, 6 and 7 occurs in human colon cancer cell lines following treatment with chloroquine (Zamame Ramirez *et al*, 2020).

Proposed mechanisms for the antiviral effect of mefloquine against coronaviruses and other viruses such as Ebola and monkeypox virus include the inhibition of viral uptake by endocytosis and plasma membrane fusion and release of the viral genome from endosomes; however, a molecular basis for the mechanism is not reported (Akazawa *et al*, 2023; Sacramento *et al*., 2022; Shionoya *et al*., 2021). In fact, MFQ’s effect on endocytosis has not been studied extensively in mammalian cells. In the malaria parasite *Plasmodium falciparum*, mefloquine inhibits the endocytosis of hemoglobin from host-erythrocytes without affecting endosome-vacuolar fusion in the parasite suggesting a specific inhibitory effect on the formation of endocytic vesicles (Hoppe *et al*, 2004). Interestingly in our studies, even at a dosing of 20 µM mefloquine, individual cells stain positive for viral proteins, suggesting that these cells internalize and replicate viral genomes (see Fig. 5B, F; Fig. 6B; Supp. Fig 4A). However, the surrounding cells do not stain positive even after being in contact with infected cells for periods of time that correspond to multiple SARS-CoV-2 replication cycles. The release/escape of coronaviruses, including SARS-CoV-2, from the cell predominantly relies on exocytosis rather than cell lysis in the first 48 h of infection (Calabrese *et al*., 2021; Cortese *et al*, 2020). To that end, coronaviruses hijack autophagic vesicles rather than exocytic vesicles for release (Chen *et al*, 2021; Miller *et al*, 2020; Münz, 2017). Mefloquine does not inhibit canonical exocytosis but is a potent inhibitor of autophagic vesicle turn-over by inhibiting lysosomal proteases such as cathepsin B (Balasubramanian *et al*, 2017; Chen *et al*., 2021; Golden *et al*, 2015; Sharma *et al*, 2012). Inhibition of lysosomal protease activity also occurs in our study. This effect varies across cell lines in the literature and was conversely interpreted in some studies as autophagy induction at concentrations that could induce a hormetic compensation (Shin *et al*, 2015; Xie *et al*, 2020). We therefore suggest that mefloquine inhibits viral release by blocking both autophagy and viral uptake.

To date, no aerosolized treatment is currently approved against coronaviruses, though local delivery to the lung to treat respiratory viruses is a logical treatment approach (He *et al*, 2022). Indeed, several groups are exploring aerosolized treatments for COVID-19, with particular interest in developing Remdesivir containing formulations (Ramsey *et al*, 2022; Vartak *et al*, 2021). However, similar to nirmatrelvir and molnupiravir, passage of SARS-CoV-2 WT-WA1 in the presence of remdesivir results in resistance mutations in the RdRp that confer resistance (i.e., 2.7-to 10.4-fold shift in EC_50_) to RDV antiviral activity (Stevens *et al*, 2022). NPs loaded with mefloquine (MFQ-NPs) exhibit a robust inhibition of coronavirus infection *in vitro* across multiple viral strains (i.e., MHV-A59, HCoV-OC43, and SARS-CoV-2 WT-WA1 and Omicron BA.1) in cells that model the human pulmonary epithelium (i.e., Calu-3). In summary, we describe a new polymeric nanoparticle system, composed of PGC-C18, which physically encapsulates hydrophobic small molecules with antiviral properties. These results encourage further *in vivo* investigation and development of MFQ-NPs for use as either a prophylactic or treatment for an array of respiratory coronaviruses.

## LIMITATIONS OF THE STUDY

Several limitations are present in the current study. First, the *in vitro* effective mefloquine concentration of ca. 17.4 µM required in our viral inhibition assays is slightly higher than in other comparable studies (Sacramento *et al*., 2022; Shionoya *et al*., 2021). This result may be a consequence of using 10-100x higher viral MOIs than comparable studies. That said, our MFQ-NPs do not release their entire drug payload in an immediate burst but rather slowly over time and exhibit low cytotoxicity and a potent antiviral effect even under a high viral burden. Our study shows that MFQ loaded NPs are an effective treatment against SARS-CoV-2 infection. More investigation is needed to determine whether MFQ-NPs release sufficient MFQ in animal lung tissues and if MFQ-NP treatment is effective in preventing or mitigating SARS-CoV-2 infection *in vivo*.

## METHODS

### Chemicals

Sodium dodecyl sulfate (L4509), chloroquine diphosphate (C6628) and mefloquine hydrochloride (M2319) were purchased from Sigma-Aldrich (St. Louis, MO). Sulfadoxine (S0899) and nitazoxanide (N1031) were purchased from TCI Chemicals (Tokyo, JP). DQ Red BSA reagent (D12051), LysoTracker Deep Red (L12492), LysoTracker Green DND-26 (L7526), and LysoSensor Yellow/Blue dextran (L22460) were purchased from Invitrogen (Waltham, MA). CellTiter 96 AQueous One Solution Cell Proliferation Assay (MTS) and CellTiter Blue were purchased from Promega (Madison, WI). Annexin V-Orange (4759) was purchased from Sartorius (Göttingen, DE) Antibodies used in this study were against the nucleoprotein of HCoV-OC43, clone 542-7D (MAB9013, Sigma-Aldrich) and against the SARS nucleocapsid protein (200-401-A50, Rockland Immunochemicals Inc, Limerick, PA).

Free base chloroquine was prepared from the phosphate salt following previously published procedures (Dodd & Bohle, 2014). Briefly, chloroquine diphosphate salt was dissolved in water in a separatory funnel and sodium hydroxide (1 M) was added until all drug precipitated. The precipitate, free base chloroquine, was extracted with dichloromethane, dried over sodium sulfate, filtered, and dried under vacuum overnight.

### Synthesis of PGC-C18 and PGC-C18-Rho

Poly(1,3-glycerol monostearate-co-ε-caprolactone) (PGC-C18) was synthesized following previously published procedures (Wolinsky et al., 2007; Wolinsky et al., 2012). Briefly, ε-caprolactone and 5-benzyloxy-1,3-dioxan-2-one monomers were combined in a Schlenk flask at a molar ratio of 4:1, respectively. This flask was then evacuated and flushed three times with N_2_. The flask was then partially submerged in an oil bath and heated to 140°C. Separately, the tin catalyst (Sn(Oct)_2_, molar ratio of monomer: initiator = 500:1) was added to a separate flask and dried under vacuum for 1 h. Dry toluene was added to the catalyst and the toluene/catalyst mixture was then injected via syringe into the monomer containing flask. The reaction was stirred at 140°C for 48 hours until the solution became viscous. The reaction was then removed from heat, and the polymer was dissolved in dichloromethane (DCM) and precipitated in cold methanol three times. The solvent was decanted, and the resulting polymer, poly(5-benzyloxy-1,3-dioxan-2-one-co-ε-caprolactone) (PGC-Bn), was dried under vacuum overnight and isolated as a white solid.

The benzyl-protecting groups were then removed via palladium-catalyzed hydrogenolysis. The resulting mixture was filtered through Celite to remove the palladium on carbon (Pd/C), yielding poly(glycerol-co-ε-caprolactone), or PGC-OH. Poly(glycerol-co-ε-caprolactone) (1 mol eq.), stearic acid (0.3 mol eq.), dicyclohexylcarbodiimide (0.24 mol eq.), and 4-dimethylaminopyridine (0.1 mol eq.) were dissolved in dichloromethane and stirred at room temperature for 18 h. The dicyclohexylurea was removed via filtration and the product, poly(1,3-glycerol monostearate-co-ε-caprolactone) (PGC-C18), was dissolved in dichloromethane (DCM) and precipitated in cold methanol three times. The solvent was decanted, and the final product, PGC-C18, was dried under vacuum overnight. Monomer and polymer structure were characterized by proton (^1^H) nuclear magnetic resonance spectroscopy (NMR) using a Varian INOVA 500 MHz instrument at the Boston University Chemical Instrumentation Center (BU-CIC). All spectra were obtained at ambient temperature with compounds dissolved in CDCl_3_ (7.25 ppm for ^1^H NMR) (Supp. Fig 1A,B). PGC-C18 polymer molecular weight and dispersity were determined against polystyrene standards using an Agilent 1260 Infinity II GPC-SEC equipped with refractive index and dual-angle light scattering detectors (Supp. Fig 1C).

For fluorescent polymer, poly(glycerol-co-ε-caprolactone) (1 mol eq.), rhodamine B (0.2 mol eq.), dicyclohexylcarbodiimide (0.22 mol eq.), and 4-dimethylaminopyridine (0.1 mol eq.) were dissolved in dichloromethane and stirred at room temperature for 18 h. Next, stearic acid (0.8 mol eq.) was added and stirred at room temperature for an additional 18 h. Lastly, the dicyclohexylurea was removed via filtration and the product, poly(glycerol monostearate-co-ε-caprolactone) rhodamine B (PGC-C18-Rho), was dissolved in dichloromethane (DCM) and precipitated in cold methanol three times. The solvent was decanted, and the final product, PGC-C18-Rho, was dried under vacuum overnight.

### Preparation of PGC-NPs

PGC-NPs were fabricated similarly to a previously described solvent evaporation approach (Ekladious *et al*., 2017). The core components, PGC-C18 (200 mg) and free drug (mefloquine, sulfadoxine, chloroquine, nitazoxanide – 25 mg, 12.5 wt%) are dissolved in 2 mL dichloromethane. This solution was placed in a sonication bath for 5 min to quickly form a homogenous solution. The surfactant, sodium dodecyl sulfate (SDS, 80 mg) is separately solubilized in 10 mM pH 7.4 phosphate buffer (8 mL). These two solutions were then added via syringe into a sonochemical reaction vessel and emulsified under an argon blanket in a pulsatile manner (30min total, 1s on / 2s off) using a Sonics Vibra-Cell VCX-600 Ultrasonic Processor (Sonics & Materials; Newtown, CT). The resulting nano-emulsion is transferred to a clean glass vial under magnetic stirring for at least 1 h to allow the dichloromethane to evaporate from the NP solution. To ensure the elimination of unassociated SDS, the nano-emulsion is then dialyzed for 24 hours against 2 L of 5 mM pH 7.4 phosphate buffer using SnakeSkin dialysis tubing (MWCO 10Kda).

Rho-NPs were fabricated similarly, with core consisting mainly of PGC-C18 (140 mg) with a small portion of fluorescent polymer, PGC-C18-Rho (60 mg). By primarily using the traditional PGC-C18 polymer, nanoparticle structure and size remains unperturbed, yet the particles fluoresce and are visible via microscopy and flow cytometry.

### Scanning electron microscopy

Unloaded NPs and MFQ-NPs were diluted 1:100 – 1:1000 times in nanopure water. Aliquots were pipetted onto silicon wafers affixed to aluminum stubs with copper tape and allowed to air dry overnight. The stubs were then sputter coated with 5 nm Au/Pd. Samples were then imaged using a Supra 55VP field emission scanning electron microscope (Carl Zeiss AG, Jena, Germany) with an accelerating voltage of 3-5 kV and working distance of 6 mm.

### Dynamic light scattering

For sizing measurements, 75 µL of NP solution is diluted in 3 mL nanopure water, and for zeta potential measurements, 30 µL of NP solution is diluted in 1.5 mL 1X phosphate buffered saline (PBS). Samples are pipetted into a cuvette and size and zeta potential are then obtained using the Brookhaven NanoBrook Omni (Brookhaven Instruments; Holtsville, NY). All measurements were performed in triplicate (n = 3).

### Quantification of drug loading and release

Small molecule drug loading (e.g., mefloquine, chloroquine, nitazoxanide, sulfadoxine) was measured using high performance liquid chromatography (HPLC). Serial dilution standards were prepared in a mobile phase composed of: 45% - 0.1% triethylamine / phosphate buffer (pH 3.0) and 55% - acetonitrile. Standard samples for each pharmacologic agent were run for 8 min through a Zorbax SB300 - C18 column (150 mm length) and detected through UV absorbance to generate a standard curve. To quantify NP loading, NPs were disrupted by adding acetonitrile to a final volume of 90% (v/v). This solution was then re-equilibrated by adding aqueous buffer to match mobile phase composition. This solution was filtered through a 0.22 µm PVDF syringe filter (Millipore) to remove large aggregates or dust prior to running samples. Samples were run similar to standards.

Mefloquine drug release from MFQ-NPs was measured using UPLC-MS (Waters ACQUITY; Milford, MA). Briefly, undiluted MFQ-NPs were loaded into Slide-A-Lyzer™ MINI Dialysis Devices, 10K MWCO (PI88401), and placed into 14 mL of release buffer (either 1X PBS (pH 7.4) with 1 v/v% Tween 20 or 0.1 M Acetate buffer (pH 5.0) with 1 v/v% Tween 20) in a conical tube. Samples were placed in a 37°C oven equipped with a shaker plate, and MFQ release was measured from the release buffer at 4, 24, 48, 72, 96, and 120 h timepoints. UPLC-MS samples were run on a Waters Acquity UPLC with a binary solvent manager, SQ mass spectrometer, Waters 2996 PDA (photodiode array) detector, and evaporative light scattering detector (ELSD) using an Acquity UPLC BEH C18 1.7 µm, 2.1 x 50 mm column (186002350). Serial dilution standards were prepared in a mobile phase composed of: 50% - release buffer (i.e., 1X PBS (pH 7.4) or 0.1 M Acetate buffer (pH 5.0)) and 50% - acetonitrile.

### qNano

Particle size and size distribution was also measured using a qNano analyzer (IZON Sciences) coupled with an adjustable nanopore (NP150), and air based variable pressure module (VPM). MFQ-PGC-NPs and carboxylated polystyrene calibration particles (CPC100, IZON Sciences) were diluted 1:500 – 1:1000 times in tris buffer electrolyte (IZON Sciences) prior to running samples according to manufacturer’s protocol. Each recorded measurement consisted of at least 500 particles counted in a 5–10 min duration. Particle size and distribution was measured using Izon control suite software.

### Nebulization of NPs

MFQ-NPs were first fabricated as described and diluted to respective concentrations using 10mM pH 7.4 phosphate buffer to prevent clogging of the vibrating meshes in the nebulizer at high concentrations. Diluted NPs were pipetted into the top reservoir of an Aerogen® Pro nebulizer and a conical tube was used to collect nebulized vapor. Centrifugation was used to condense the vapor into a liquid solution. Nebulized MFQ-NPs were compared to the pre-diluted sample as well as pre-nebulized dilutions for size and morphological changes using DLS and SEM as described.

### Cell culture

Vero E6 cells were obtained from the American Type Culture Collection (CRL-1586, ATCC, Manassas, Virginia) and grown in Eagle’s Minimal Essential Medium (Corning, Corning, NY) supplemented with 10% FBS and Penicillin (100 U/mL)/Streptomycin (100 μg/mL) (PenStrep, Gibco, Waltham, MA). Calu-3 human lung cancer cells were obtained from ATCC (HTB-55) and grown in EMEM supplemented with 10% FBS and PenStrep. L929 and 17CL-1 mouse fibroblast cells were a kind gift from Volker Thiel’s lab at the University of Bern and Elke Mühlberger’s lab at Boston University and grown in EMEM supplemented with 10% FBS and Primocin (100 μg/ml) (Invivogen, San Diego, CA). HCT-8 cells were obtained from ATCC (CCL-244) and grown in RPMI-1640 (ATCC) supplemented with 10% horse serum. HFL1 cells were obtained from ATCC (CCL-153) and grown in F-12K media supplemented with 10% FBS and PenStrep.

### Viruses

SARS-CoV-2 USA-WA1/2020 (referred to as WT-WA1) and Omicron BA.1 were obtained from the Biodefense and Emerging Infections (BEI) Resources of the National Institute of Health (contributed by Mehul Suthar). All work with SARS-CoV-2 was performed at the UCLA high containment laboratory at biosafety level 3. SARS-CoV-2 was propagated and passaged in Vero E6 cells. HCoV-OC43 was obtained from ATCC (VR-1558) and propagated in HCT-8 cells. MHV-GFP was a kind gift from Volker Thiel’s lab at the University of Bern and Elke Mühlberger’s lab at Boston University. MHV-A59-GFP was propagated in 17CL-1 cells. Viral titers were determined by assessing viral cytopathic effect (CPE) by microscopy in cells infected with serial 10-fold dilutions respectively. TCID_50_/ml was calculated using the Reed-Muench method.

### *In vitro* cell viability

The cytotoxicity of NPs with and without MFQ was evaluated using a tetrazolium-based MTS cell proliferation assay (Promega CellTiter 96 Aqueous One Solution Cell Proliferation Assay). HFL1, Calu-3, and Vero E6 were cultured in a 96-well plate at 12,000 cells/well for 1 day, after which the media was exchanged for media containing no treatment, unloaded PGC-NPs (1.95, 3.91, 7.81, 15.63, 31.25, 62.5, 125, 250, 500, 1,000 µg/mL NPs), MFQ-NPs (1.95, 3.91, 7.81, 15.63, 31.25, 62.5, 125, 250, 500, 1,000 µg/mL NPs), free MFQ solubilized in DMSO (0.78, 1.56, 3.13, 6.25, 12.5, 25, 50 µM), or DMSO as a vehicle control (0.031, 0.063, 0.125 % v/v relative to culture media). The cells were then incubated with treatment for 24 hours, after which cell viability was quantified with a SpectraMax iD3 (Molecular Devices) plate reader relative to the no treatment control, after correcting for background absorbance. Alternatively, toxicity was determined using the CellTiter Blue assay (Promega). Vero E6, or Calu-3 cells were seeded in clear 96-well plates and treated with MFQ-NPs at 1 μg/mL – 800μg/mL, unloaded NPs, or equivalent concentrations of free MFQ solubilized in dimethyl sulfoxide (DMSO) for 72 h. Subsequently, CellTiter reagent was added to the wells for 1 h and fluorescence emission was measured at ex:565 nm, em:620 nm using a Tecan Spark 10M plate reader (Tecan Ltd., Männedorf, Switzerland).

### Cellular uptake of Rho-PGC-NPs via flow cytometry

FACS analysis of Rho-NPs treated cells was performed with an Attune NxT Flow Cytometer (Invitrogen). HFL1, Calu-3, and Vero E6 were cultured in a 96-well plate at 12,000 cells/well for 1 day, after which the media was exchanged for media containing 75 µg/mL Rho-NPs. The cells were then incubated with treatment for 24 hours, after which cells were trypsinized, washed with PBS by centrifugation, resuspended in FACS buffer (PBS + 2% FBS) and then subjected to flow cytometry. Cell debris was excluded by gating on the forward and side scatter plot. Intensity bar graph for uptake is displayed as mean values calculated from median fluorescence intensity for cell population identified as positive based on the gating strategy.

### Cellular localization of Rho-PGC-NPs via confocal microscopy

HFL1 were grown on 12-well glass bottom plates (Celvis, P12-1.5H-N) at a density of 50,000 cells/well and grown for 24 h. Cells were then treated with a 75 µg/mL dose of Rho-NPs for 1 h, 4 h, or 24 h. Cell were then washed with 1X PBS and incubated with 50 nM LysoTracker Deep Red (Invitrogen) in culture media for 1.5 h. Following acidic organelle labelling, cells were washed with 1X PBS and incubated with 1 µg/mL Hoechst 33342 (Thermo) and 5 μg/mL Wheat Germ Agglutinin Oregon Green 488 Conjugate (Invitrogen) for 10 min at 37 °C. Cells were then washed with 1X PBS and incubated with 1 µg/mL Hoechst 33342 in Live Cell Imaging Solution (Thermo) and imaged immediately on an Olympus FV3000 confocal microscope with a temperature control chamber at 37 °C. Cells were imaged using a 60X oil immersion objective. Co-localization of Rho-NPs and Lysotracker dye was determined by calculating Pearson’s coefficient using CellProfiler (Carpenter *et al*, 2006).

### Lysosensor-Dextran Yellow/Blue imaging

Vero E6 cells were seeded at a density of 10,000 cells/well into Greiner CellView 4 compartment dishes. After 24 h cells were stained with 5 mg/mL Lysosensor-Dextran Yellow/Blue dye in EMEM for 3 h following an overnight media chase. The next day cells were incubated for 24 h with 100 µg/mL NPs (±MFQ) or 10 or 20 µM MFQ. Bafilomycin A1 was added at a concentration of 200 nM as alkalization agent 2-4 h before imaging. Imaging was performed using a Zeiss LSM880 (Carl Zeiss AG, Jena, Germany) equipped with a Coherent 2-photon laser at a 2-photon excitation of 720 nm and 2 emission bands at 400-480 nm (blue) and 510-620 nm (yellow) using a 63X oil immersion objective. pH standard curves were generated by permeabilizing Lysosensor-Dextran Yellow/Blue stained cells with 10 µM nigericin, 20 µM monensin in pH clamped buffers ranging from pH 4.5 – 6.0. Lysosomal ROIs and Yellow/Blue staining intensity were determined using CellProfiler.

### Lysotracker Green imaging

Vero E6 cells were seeded at a density of 10,000 cells/well into 96-well Greiner µClear imaging plates. After 24 h cells were treated with 50-100 µg/mL NPs (±MFQ), 20 µM MFQ, or 200 nM bafilomycin A1 for 24 h and subsequently stained with 1 µM Lysotracker Green and Hoechst 33342. Cells were imaged using an Operetta High-Content Imager (PerkinElmer Inc, Waltham, MA) using a 20X air objective and lysosomal accumulation defined as Lysotracker positive signal area % in expanded nuclear ROIs. Images were analyzed using CellProfiler.

### DQ-Red BSA assay

HFL1 cells were cultured in a 96-well plate at 15,000 cells/well for 24 h, after which the media was exchanged for media containing no treatment, empty NPs (12.5, 25, 50, 75, 100 µg/mL NPs) or MFQ-NPs (12.5, 25, 50, 75, 100 µg/mL NPs) for 24 h. Control treatments include Bafilomycin A1 (200 nM), and Pepstatin A (10 µg/mL) + E64d (10 µg/mL) for 4 h, or free MFQ (10, 15 µM) for 24 h. Following treatment, cells were washed with 1X PBS and incubated with 10 µg/mL DQ Red BSA reagent (Thermo) in culture media for 1 h. After incubation with assay reagent, cells were trypsinized, washed with 1X PBS by centrifugation, resuspended in FACS buffer (PBS + 2% FBS) and then subjected to flow cytometry. Cell debris was excluded by gating on the forward and side scatter plot. Relative protease activity is displayed as mean values calculated from median fluorescence intensity for cell population identified as positive based on the gating strategy.

### MHV-GFP infection kinetics determination

In order to determine the best timepoint to assess MHV-A49 (MHV-GFP) infection. L929 cells were seeded in clear bottom Corning 96-well plates and inoculated with MHV-GFP at an MOI of 0.1. A 1:500 dilution of Annexin V-Orange (Sartorius) was added to the cells to monitor MHV induced cell lysis and cell death. Cells were imaged every hour for 48 h using an OmniFL live cell analysis platform (CytoSMART Technologies, Eindhoven, NL) and growth curves were determined using Cytosmart cloud software.

### NP treatments and viral infections

For MHV-A49 infection and HCoV-OC43 assays L929 or Vero E6 cells are plated into 96-well Corning imaging plates. 24 h later, cells undergo a 1 h prophylactic pre-treatment with NPs ± MFQ (12.5-100 µg/mL), free MFQ (1.25-20 µM), positive control treatment (10 μM remdesivir), or no treatment. Following pre-treatment, media is exchanged with serum free media containing virus (HCoV-OC43 at MOI of 1 or MHV at MOI of 0.1) for 1 h. After 1 h of inoculation, viral containing media is removed, wells are rinsed with 1X PBS and treatments are added back to the wells for another 24-48 hrs. Cells are then fixed with 4% paraformaldehyde, permeabilized with 0.1% Triton X-100 in PBS, blocked with 2% BSA, 5% normal donkey serum (NDS), and infection is visualized by immunofluorescence staining of pan-corona-nucleocapsid protein for HCoV-OC43 with a mouse monoclonal primary and a donkey-anti-mouse AlexaFluor488 conjugated secondary antibody and cell nuclei counterstaining with DAPI. MHV-GFP positive cells will appear GFP-fluorescence positive or as syncytia.

For SARS-CoV-2 infection assays, Vero E6 or Calu-3 cells were plated into 96-well, clear-bottom imaging plates. 24 h later, cells undergo a 1 h prophylactic pre-treatment in 100 µL of media containing 2X concentrated NPs ± MFQ (25-200 µg/mL), free MFQ (2.5-40 µM), positive control treatment (20 μM remdesivir), or no treatment. Following pre-treatment, 100 µL of media containing virus (SARS-CoV-2 at MOI of 0.1 for Vero E6 and MOI of 0.2 for Calu-3 cells) is added to the wells diluting the treatments to 1X for another 48 h. Alternatively, for post-inoculation treatments, Calu-3 cells are infected with SARS-CoV-2 MOI 0.2 diluted in serum free EMEM for 1 h. Subsequently, the inoculum is removed, the cells washed once with DPBS, and treated with NPs ± MFQ (12.5-100 µg/mL), free MFQ (1.25-20 µM), positive control treatment (10 μM remdesivir), or no treatment for 48 h. Cells are then fixed with 4% paraformaldehyde, permeabilized with 0.1% Triton X-100 in PBS, blocked with 2% BSA, 5% NDS, and infection is visualized by immunofluorescence staining of SARS-CoV-2 N protein with a rabbit polyclonal primary and a donkey-anti-rabbit AlexaFluor568 conjugated secondary antibody and cell nuclei counterstaining with DAPI. Plates were imaged with an Operetta High-Content imager using a 10X air objective, and images were processed using CellProfiler. Positive cells were determined as expanded nucleus ROIs containing above threshold virus positive staining, syncytia were defined as irregularly close nucleus clusters of >40 µm diameter.

### Expression of viral entry proteins

Calu-3 or L929 cells were plated into 12-well plates at 130,000 or 100,000 cells/well, respectively. 24 h later, the media was exchanged for media containing no treatment, unloaded PGC-NPs (50, 100 µg/mL NPs), MFQ-NPs (50, 100 µg/mL NPs), free MFQ solubilized in DMSO (8.7, 17.4 µM), or DMSO as a vehicle control (0.044, 0.087 % v/v relative to culture media). The cells were then incubated with treatment for 48 h, after which RNA was isolated using the RNeasy Plus Mini Kit (Qiagen) according to manufacturer’s protocol. Total RNA concentration was determined using Nanodrop (Thermofisher Scientific) and cDNA was generated using the High-Capacity cDNA Reverse Transcription Kit (4368813, Applied Biosystems). TaqMan probes (ThermoFisher) were used to measure expression level of membrane bound proteins responsible for viral uptake or processing enzymes essential for the production of said transmembrane proteins: *hACE2* and *hTMPRSS2* in Calu-3 cells. Additionally, we measured the expression level of Cathepsin L (*hCTSL*) which is a lysosomal protease essential for S protein processing and endolysosomal membrane fusion in the SARS-CoV-2 endocytic infection route. Similarly, TaqMan probes were used to measure expression level of *mCEACAM1* in L929 samples.

Changes in relative gene expression were quantified using the 2^-ΔΔCT^ method and *hGAPDH* or *mGAPDH* were used as housekeeping genes for Calu-3 and L929, respectively.

### SARS-CoV-2 uptake analysis by RT-qPCR

For the SARS-CoV-2 uptake analysis, Vero-E6 cells were seeded into 6-well plates at densities of 5×10⁵ cells/well and incubated overnight at 37°C with 5% CO₂. The next day, cells were pre-treated with varying concentrations of nanoparticles (NPs) or compounds for 1 hour in serum-free media (SFM). Following pre-treatment, cells were exposed to SARS-CoV-2 for 1 h at a multiplicity of infection (MOI) of 0.2 at 4°C (to assess adherence) or at 37°C (to assess endocytosis and replication) in the presence of the respective compounds and NPs in SFM. For adherence-only samples, cells were washed with ice-cold PBS immediately after the infection period and scraped into 500 µL of TRI reagent (T9424, Sigma). For endocytosis and replication samples, cells were washed with warm PBS after the infection period and then incubated in growth media containing the compounds and NPs for 6 and 24 hours, respectively, before collection in TRI reagent as previously described. Samples were stored at -80°C until further processing for RT-qPCR.

RNA extraction from cell homogenates was performed using TRI Reagent according to the manufacturer’s guidelines. The reverse transcription of RNA to cDNA was carried out using the High-Capacity cDNA Reverse Transcription Kit (4368813, Applied Biosystems), following the manufacturer’s instructions. The reverse transcription protocol included an initial 10-min incubation at 25°C, followed by 120 min at 37°C, 5 min at 85°C, and indefinite maintenance at 4°C.

Quantitative PCR (qPCR) was conducted using the PowerSYBR Green PCR Master Mix (4367659, Thermo), adhering to the manufacturer’s protocol. The qPCR cycling conditions were an initial 2-minute incubation at 50°C, 10 minutes at 95°C, followed by 40 cycles of 15 seconds at 95°C and 60 seconds at 60°C, with SYBR wavelength acquisition. The temperature ramp rate for all cycling steps was set at 1.6°C/second. Each qPCR run included triplicate technical replicates, a 2019-nCOV positive control plasmid (10006621, IDT), and an infection-negative control sample. Data analysis was performed using the ΔΔCt method for fold change calculation.

### Statistical analysis

Unless otherwise noted, all experiments were repeated at least three times. Unless otherwise mentioned data were graphed and analyzed using GraphPad Prism 9.1. Data are displayed as means ±SD or SEM where appropriate. To determine statistically significant differences one-way and two-way ANOVA analysis with Dunnet post-hoc tests were applied where appropriate.

## ACKNOWLEDGEMENTS

The authors would like to thank Drs. Stan Louie and Daniel Dagan for valuable discussion and critical reading of the manuscript. We would also like to thank the team at Aerogen (Patricia Dailey, Ronan MacLoughlin, and David Coyle) for gifting multiple Aerogen® Solos and controllers for nebulization. In addition, we would like to thank Dr. David Ferrick and the team at Axion Biosystems for the kind gift of the CytoSMART OmniFL. This project was supported by UCLA COVID-19 Emergency Response OCRC Grant #20-21 to Orian Shirihai. B.M.T. and A.M. were funded through the BUnano Fellowship Program.

## AUTHOR CONTRIBUTIONS

Conceptualization: A.P., B.M.T., A.H.C., M.W.G., and O.S.S

Methodology: A.P., B.M.T., and V.A.

Advice: V.A.

Investigation: A.P., B.M.T., A.M., S.A., A.J.B, R.A.S, B.S., G.G. Jr., and M.V.

Visualization: A.P., B.M.T., A.M., S.A.

Supervision: A.H.C., M.W. G., and O.S.S

Writing—original draft: A.P., B.M.T.

Writing—review & editing: All authors

Funding: B.M.T., A.M., M.W. G., and O.S.S

### Data Availability

The raw data required to reproduce these findings are available from the authors upon request.

## EMAILS

Brett M. Tingley

Anton Petcherski

Andrew Martin

Sarah Adams

Alexandra J. Brownstein

Ross A. Steinberg

Byourak Shabane

Corey Osto

Jennifer Ngo

Gustavo Garcia Jr.

Michaela Veliova

Vaithilingaraja Arumugaswami

Aaron H. Colby

Orian S. Shirihai

Mark W. Grinstaff

## FIGURE LEGENDS

**Supplemental Figure 1.**
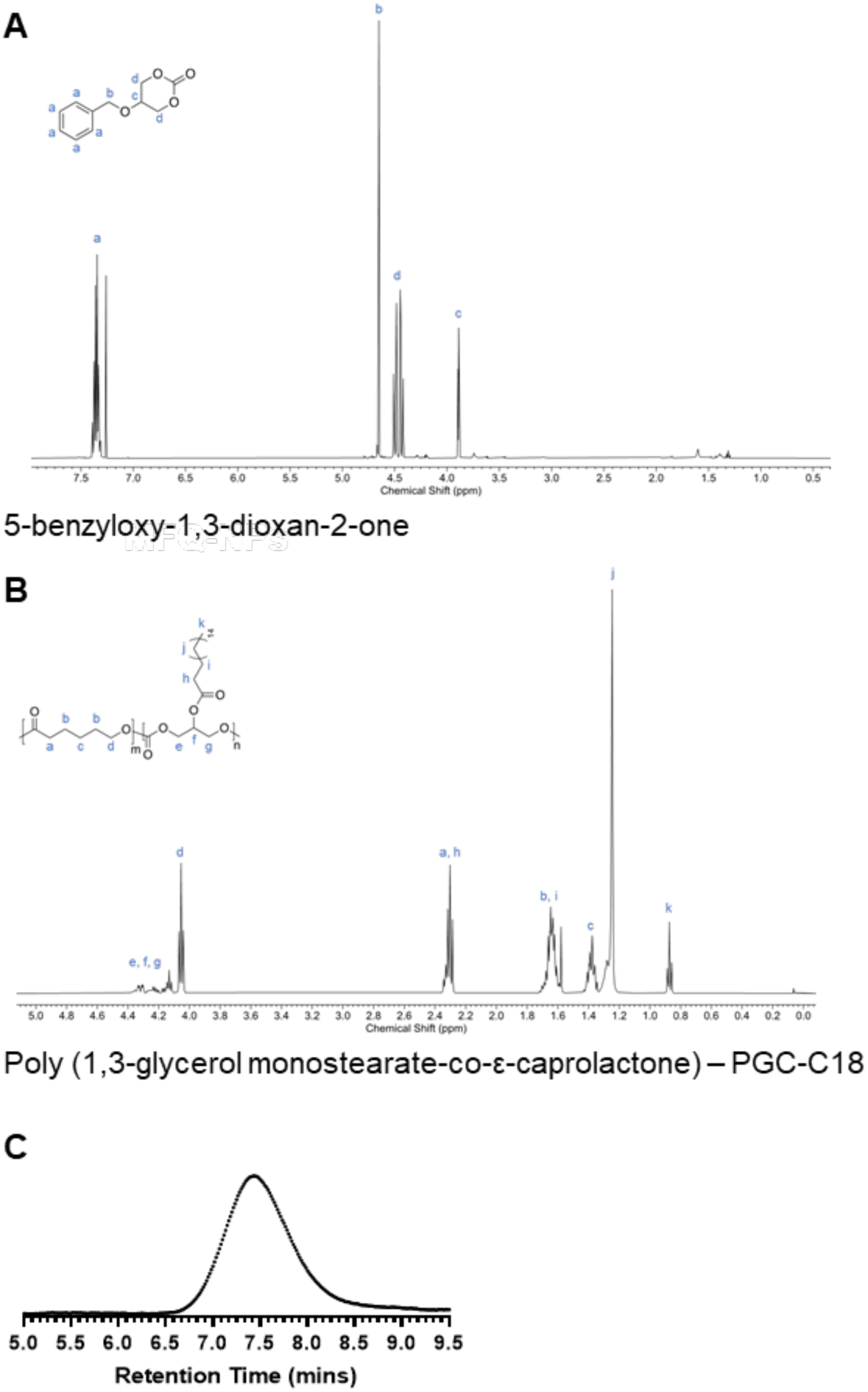
(A) ^1^H NMR (500 MHz, CDCl_3_) spectrum of 5-benzyloxy-1,3-dioxan-2-one monomer. (B) ^1^H NMR (500 MHz, CDCl_3_) spectrum of poly(glycerol monostearate-co-ε-caprolactone) (PGC-C18) using a 4:1 molar ratio of ε-caprolactone to 5-benzyloxy-1,3-dioxan-2-one monomers. (C) THF GPC trace of PGC-C18. PGC-C18 molecular weight (M_n_) = 78272 g/mol and dispersity (Đ) = 1.668 were determined based on polystyrene standards according to refractive index (RI) detector.

**Supplemental Figure 2.**
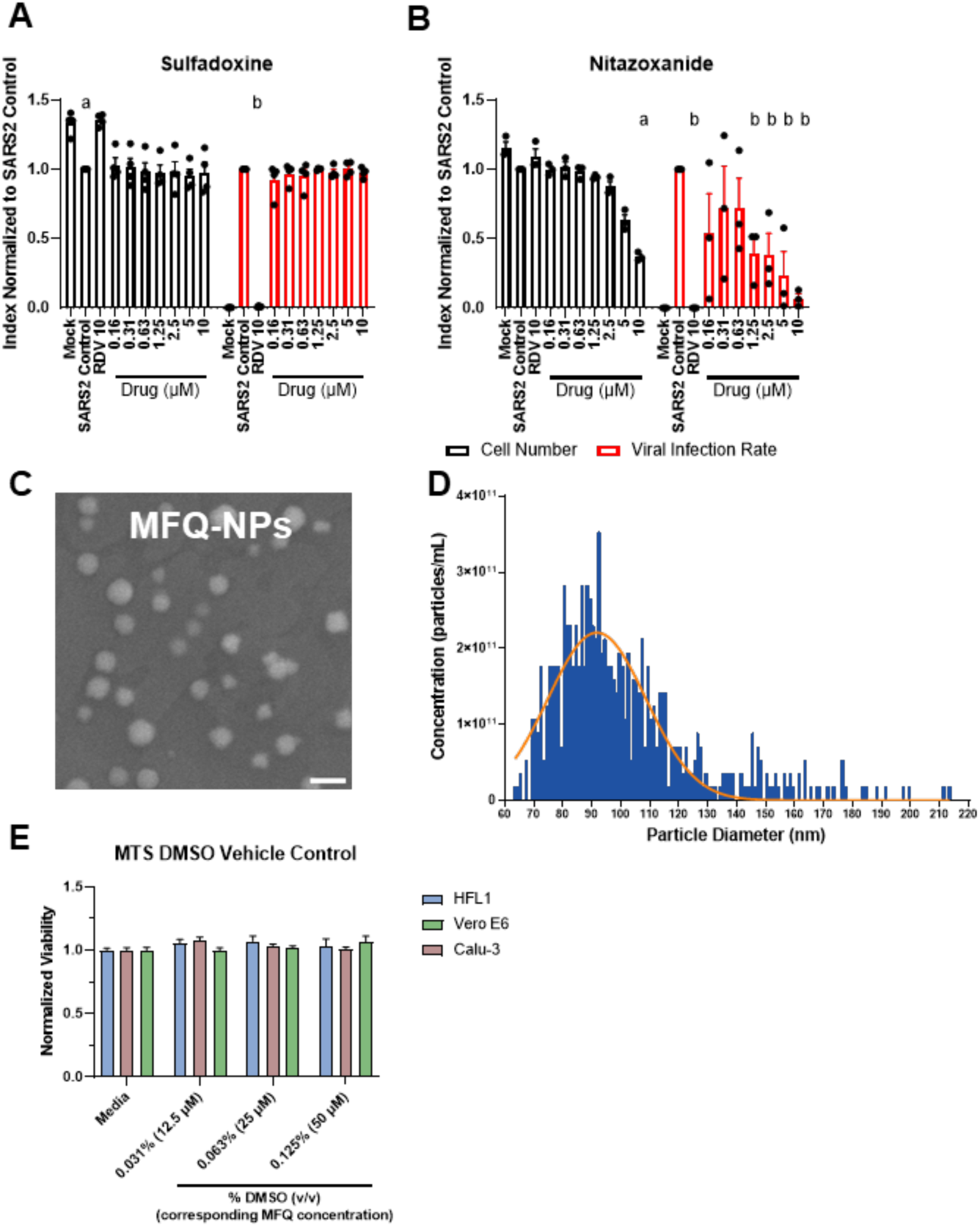
(Connected to Figures 1 and 2). (A) Quantification of cell numbers and fractions of infected cells in free molecular sulfadoxine pretreated Vero E6 cells infected with SARS-CoV-2 WT-WA1. (B) Quantification of cell numbers and fractions of infected cells in free molecular nitazoxanide pretreated Vero E6 cells infected with SARS-CoV-2 WT-WA1. (C) Electron micrograph of MFQ-NPs demonstrates consistent size and morphology. Scale Bar=200 nm. (D) Size distribution of MFQ-NPs measured by tunable resistive pulse sensing (i.e., qNano). (E) Effects of different concentrations of DMSO-vehicle treatments on HFL1, Vero E6, and Calu-3 cell viability measured by MTS assay. All biological replicates represent the mean of n=3 technical replicates. All experiments represent N=3-4 biological replicates. Sizing and dispersity data are displayed as means ±SD; all other data are displayed as means ± SEM.

**Supplemental Figure 3.**
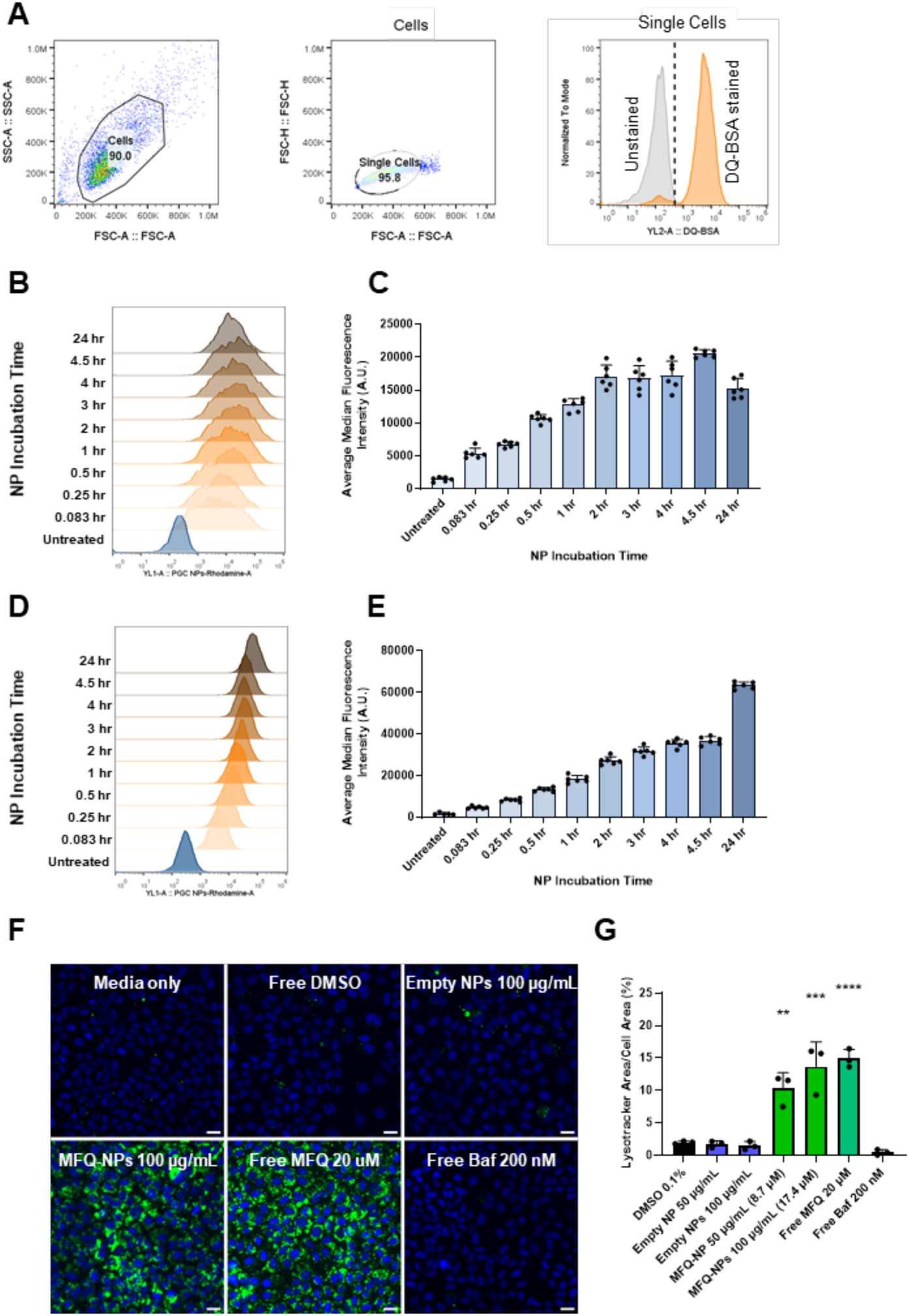
(Connected to Figures 3 and 4). (A) General pipeline outlining FlowJo gating of viable, single cells using forward and side scattering, resulting in histogram frequency distribution of individual cell fluorescence. Positive cell gating is based on unstained and untreated controls, representative unstained and stained controls from the DQ-Red BSA assay are shown. Rho-NP uptake measured by flow cytometry in Vero E6 cells and displayed as (B) representative intensity histogram or (C) median intensity bar graph. Rho-NP uptake measured by flow cytometry in HFL1 cells and displayed as (D) representative intensity histogram or (E) median intensity bar graph. (F) Representative images of Vero E6 cells treated with NPs ± MFQ (100 µg/mL), free MFQ (20 µM), bafilomycin A1 (200 nM), or control and stained with Hoechst 33342 (blue) and Lysotracker Green (green) to probe for lysosome accumulation. Scale bars=20 μm. (G) Quantification of lysosomal accumulation. Statistical significance was determined by one-way ANOVA: **p<0.01, ***p<0.001, ****p<0.0001 against untreated controls. All experiments represent N=3 biological replicates which represent the mean of at least n=3 technical replicates. Intensity histogram displays a single representative cell population per timepoint. Intensity bar graph is displayed as medians ±SEM. All other experiments are displayed as means ± SEM.

**Supplemental Figure 4.**
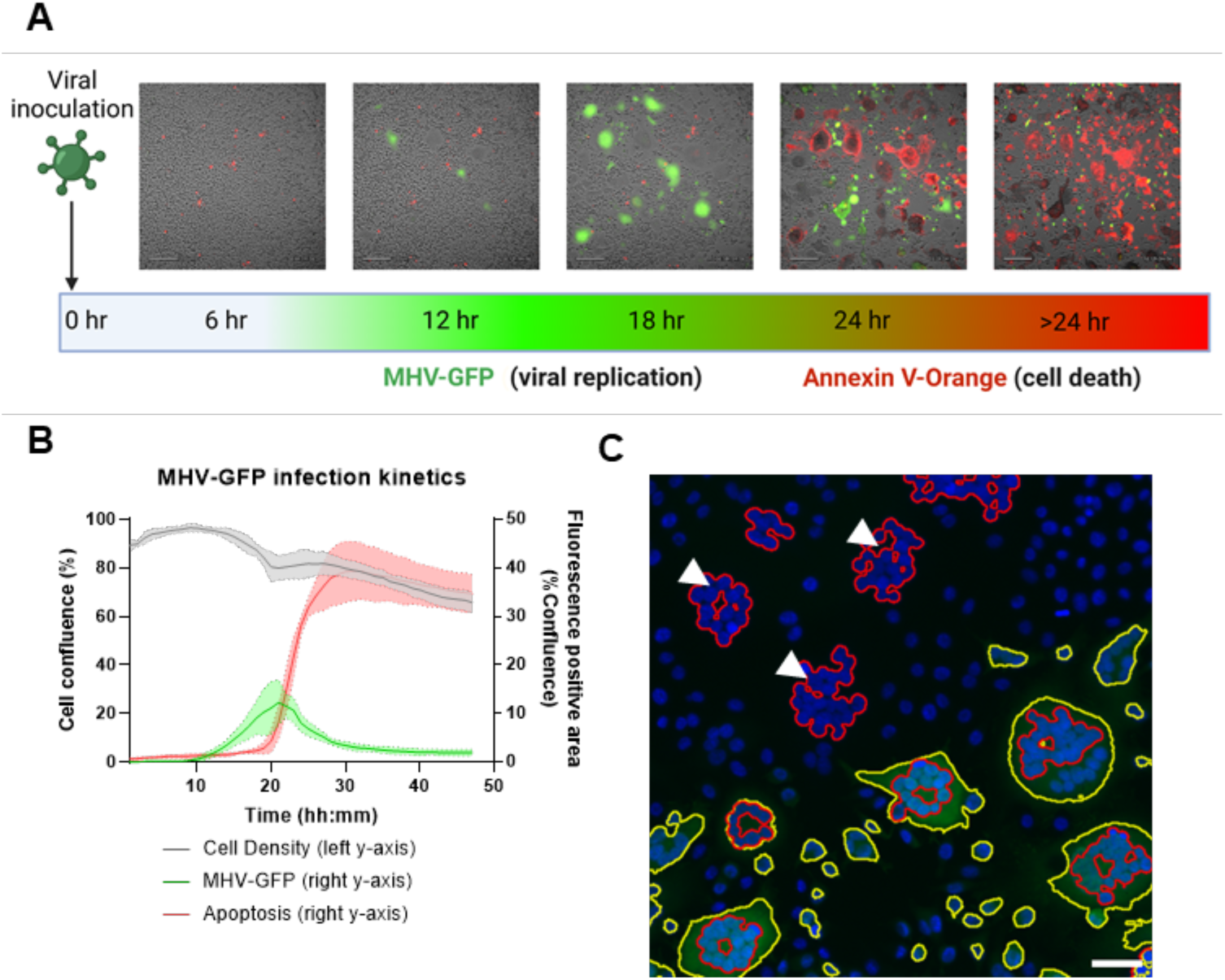
(Connected to Figure 5). (A) Timeline of viral inoculation, GFP expression, and apoptosis in MHV-GFP infected L929 cells obtained by continuous image-based monitoring with a Cytosmart OmniFL analysis platform. Brightfield cell images are overlayed with GFP fluorescence (green) and Annexin V-Orange fluorescence (red). Scale bars=200 µm. (B) Representative viral infection kinetic quantification over a 48 h infection period. (C) Representative example of syncytia (red outlines) recognized by CellProfiler analysis as clustered nuclei and GFP-positive area (yellow outlines) in fluorescence microscopy images obtained with the Operetta high-content imager. Nuclei (blue) are stained with DAPI, MHV (green) is visualized by GFP expression. GFP-negative syncytia are marked with white arrowheads. Scale bar=50 µm. Cytosmart Omni FL experiment biological replicates represent the mean of n=3 technical replicates. Experiments represent N=3 biological replicates.

**Supplemental Figure 5.**
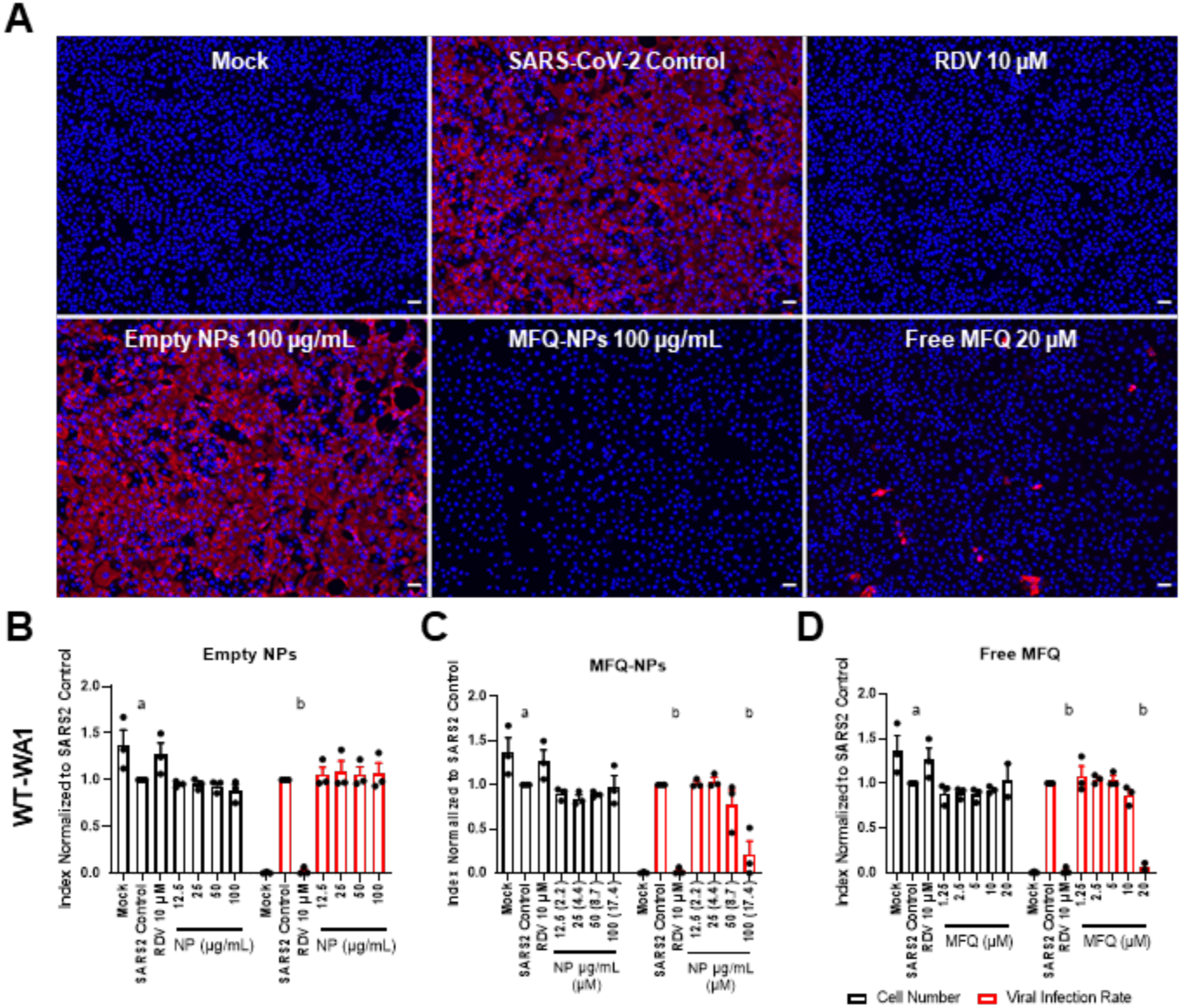
(Connected to Figure 6). (A) Schematic describing the treatment and infection sequence of prophylactic NP treated SARS-CoV-2 infected cells. (B) Representative images of Vero E6 cells pre-treated with control, RDV (10 µM), free MFQ (20 µM), NPs ± MFQ (100 µg/mL) and infected with SARS-CoV-2 WT-WA1 (MOI 0.1) for 48 h. Nuclei were stained with DAPI (blue) and virus infected cells visualized by immunofluorescence staining against SARS-CoV-2 N-protein (red). Scale bars=50 μm. Quantification of cell numbers and fractions of infected cells in unloaded PGC-NP (C), MFQ-NP (D) or free MFQ (E) pretreated Vero E6 cells infected with SARS-CoV-2 WT-WA1. Statistical significance was determined by two-way ANOVA: a (at least) p<0.05 against SARS-CoV-2 Control cell numbers; b (at least) p<0.05 against SARS-CoV-2-Control infected cell area. All biological replicates represent the mean of n=3 technical replicates. All experiments represent N=3 biological replicates. Data are displayed as means ± SEM.

**Supplemental Figure 6.**
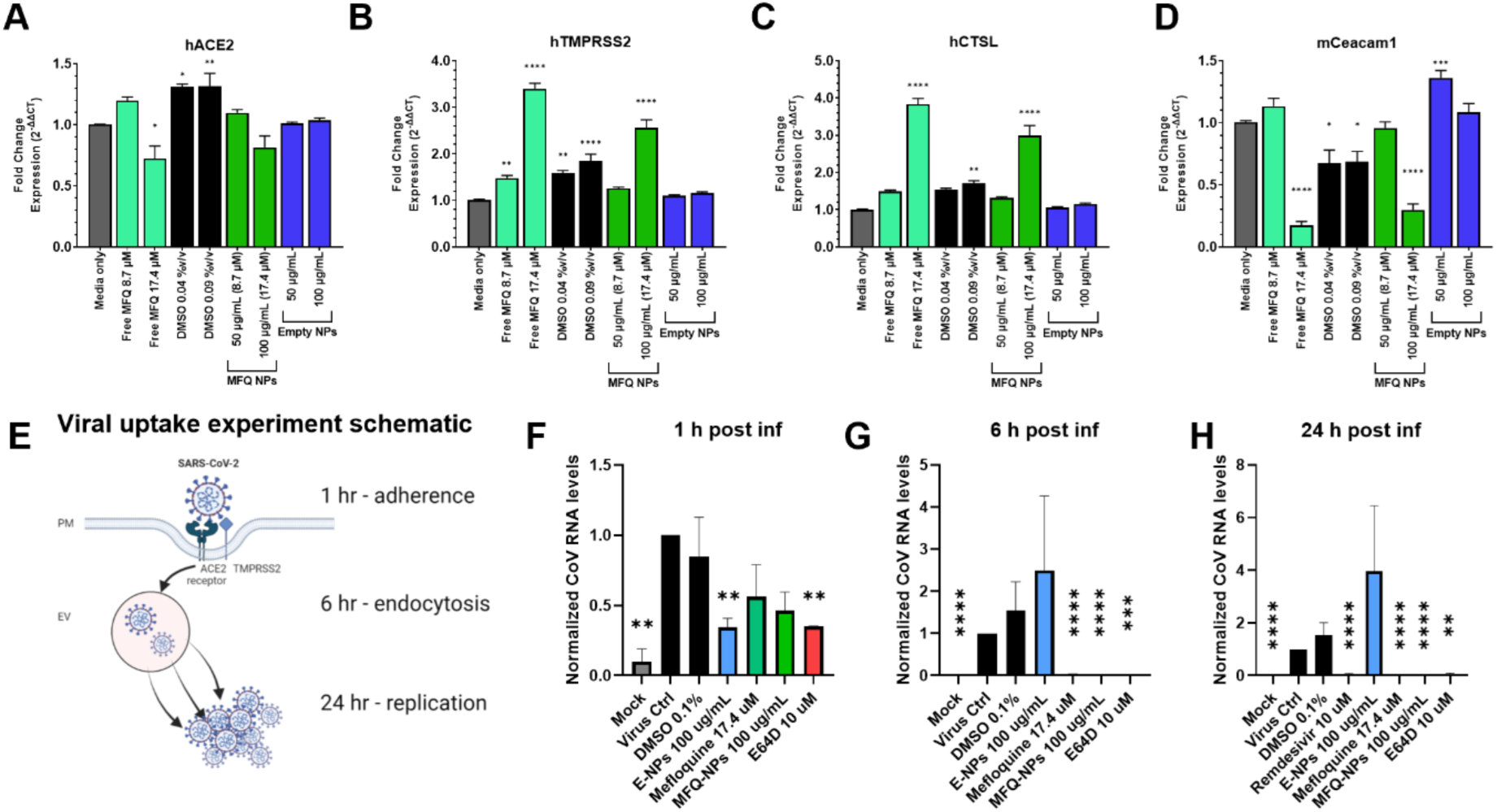
RNA expression level of essential proteins responsible for viral uptake and temporal NP effects on viral adherence and uptake. (A) ACE2 expression in Calu-3 cells, (B) TMPRSS2 expression in Calu-3 cells, (C) Cathepsin L expression in Calu-3, and (D) CEACAM1 expression in L929 cells following 48 h treatment with unloaded PGC-NPs, MFQ-NPs, free MFQ solubilized in DMSO, DMSO as a vehicle control, or no treatment. (E) Schematic representation of the experimental timeline of SARS-CoV-2 uptake into the cell. Viral RNA levels after (F) 1 h, (G) 6 h, and (H) 24 h of viral incubation in Vero cells pre-treated with empty PGC-NPs, MFQ-NPs, free MFQ solubilized in DMSO, DMSO as a vehicle control, or no treatment. All biological replicates represent the mean of n=3 technical replicates. All experiments represent N=3-5 biological replicates using 3 independent PGC-NP or MFQ-NP batches. Data are displayed as means ± SD (A-D) or SEM (F-H). Statistical significance was determined by one-way ANOVA: *p<0.05, **p<0.01, ***p<0.001, ****p<0.0001 against untreated controls or a mixed-effects ANOVA: *p<0.05, **p<0.01, ***p<0.001, ****p<0.0001 against untreated controls.

